# Calcium homeostatic feedback control predicts atrial fibrillation initiation, remodeling, and progression

**DOI:** 10.1101/2024.10.24.619898

**Authors:** Nicolae Moise, Seth H. Weinberg

## Abstract

Atrial fibrillation (AF) is a progressive disorder, with arrhythmia episodes becoming increasingly longer and ultimately permanent. The chaotic electrical activity by itself is well-known to drive progression, a process classically summarized as “AF begets AF.” However, the mechanisms underlying this progression are not yet well defined. We hypothesize that calcium homeostatic feedback regulating ion channel expression is a critical mechanistic component of this pathological process. We propose a modeling framework that tracks both short-term beat-to-beat electrical and calcium activity and long-term tissue substrate remodeling as a single coupled dynamical system. Importantly, the full AF progression from healthy to pathological remodeled tissue is reproduced, in contrast with prior studies that consider “snapshots” of various AF stages. Simulations predict that single cells respond to fast pacing by maintaining intracellular calcium concentrations through dynamic ion channel expression and electrical phenotype changes. In two-dimensional (2D) homogeneous tissue, spontaneous spiral waves stabilize into permanent re-entry. In 2D heterogeneous tissue, we observe the initiation of re-entrant activity in response to fast pacing, followed by increasingly longer intermittent, and then permanent, arrhythmic activity. Simulations predict critical properties of re-entrant wave locations, leading to a novel hypothesis: spiral wave activity itself drives underlying substrate remodeling and the emergence of remodeled tissue “niches” that support the stabilization of fast re-entrant activity. Thus, the model joins multiple lines of inquiry (i.e., long-term calcium regulation, ion channel co-expression and remodeling, and tissue-scale arrhythmia spatiotemporal organization) into a single coherent framework, and for the first time, captures the dynamics of the long-term natural history of AF.

## 1 Introduction

Atrial fibrillation (AF) is the most common cardiac arrhythmia and the leading cause of stroke, causing significant morbidity and mortality [1]. AF is characterized by continuous, chaotic electrical waves that prevent regular atrial activation and contractile function. AF is noteworthy for its natural history: sporadic, self-terminating episodes of disordered activity (known as paroxysmal AF) become more and more frequent, ultimately leading to a state of continuous chaotic activity, or permanent AF.

In a process that has been summarized succinctly as “AF begets AF” [2], the electrical activity itself directly causes changes in the underlying atrial tissue substrate. Specifically, rapid electrical activity drives electrical remodeling: decreased inward sodium current (*I*_*Na*_) and calcium current (*I*_*CaL*_), and increased inward-rectifier potassium current (*I*_*K*1_) and sodium-calcium exchanger (*I*_*NCX*_) among the most critical effects [3]. These changes in ion channel expression collectively determine a shorter atrial action potential duration (APD), which in turn increase the likelihood of re-entrant electrical activity. Longer lasting (chronic) AF further leads to structural remodeling consisting of heterogeneous gap junction coupling and fibrosis, which in turn slow conduction, and amplify the pro-arrhythmic effects of electrical remodeling [4, 5].

Critically, all of these changes lead to greater likelihood of AF initiation and persistence, thus creating a vicious cycle. However, due to the intrinsically long timescales associated with this cycle, studies almost exclusively focus on the beginning and the end of this process, comparing properties of healthy atrial tissue with tissue following AF-associated remodeling. Computational studies similarly simulate these binary conditions based on either healthy or AF-associated model parameters. The goal of this study is to reproduce and understand the *dynamics*, not just *endpoint*, of the progressive nature of AF, which is inherently a spatio-temporally complex process that depends on the coupled history and organization of arrhythmic events and dynamic atrial tissue remodeling.

Regulation of ion channel expression has been extensively studied in excitable cell systems, such as the neuron. O’Leary and colleagues [6] proposed an elegant mathematical model, in which neuronal ion channel expression levels are governed by a feedback system maintaining a particular homeostatic intracellular calcium (*Ca*_*i*_) “set point” or “target” (*Ca*_*tgt*_) and correlations between the different channels, thus ensuring physiological cellular electrical characteristics. We recently adapted this feedback model to describe the initiation of pacemaking activity in the sinoatrial node in the heart [7]. Importantly, we identified that, while ion channels are intrinsically regulated within an individual myocyte, electrical activity mediates interactions between myocytes (i.e., tissue-scale dynamics), thus producing emergent patterns for the spatial organization of pacemaking activity. Critically, the feedback model captures the inherent multiple timescales of these processes: millisecond-to-second scale dynamics of ion channel gating and fluxes and and hours-todays scale dynamics of ion channel expression remodeling.

We hypothesize that the same theoretical framework can be expanded to explain the initiation and progression of AF in atrial tissue. The two key properties of the feedback system described above, correlated ion channel expression and dynamic expression regulated by *Ca*_*i*_, are also found in the heart. First, ion channels are expressed in a coordinated way in the heart [8]: it has been shown that Kir2.1 and Nav1.5 proteins reciprocally increase each others’ expression [9] and are trafficked together to the cell membrane [10]. Further, a number of channel mRNAs (CACNA1C, SCN5A, KCNQ1, hERG1a) are co-localized and co-transcribed [11]. These mechanisms lead to a balance of depolarizing and repolarizing currents [12], which has also been demonstrated computationally by optimizing for ‘good enough solutions’ that match APD and calcium transient (CaT) characteristics [13]. Second, rapid pacing is well-known to drive changes in ion channel expression characteristic of AF, and there is abundant evidence that calcium, and specifically pacing-induced *Ca*_*i*_ overload, is the main driver [5]. AF-associated changes ultimately converge on reducing CaT amplitude, as well as reduced APD, and thus decrease average *Ca*_*i*_. Notably, blocking excessive *Ca*_*i*_ entry using calcium channel blockers can prevent remodeling [14, 15]. Finally, remodeling induced by pacing occurs in a variety of species (e.g., sheep [16], dog [17], goat [2], horse [18]) and across cell types (e.g. cardiac iPSCs [19], isolated atrial [20] and ventricular cells [21]), suggesting that this mechanism is a general homeostatic response to calcium overload.

Here, we are interested in answering the question: how does AF initiate and progress? Current computational efforts have focused on increasingly more spatial detail, from whole atria models [22] to subcellular structures [23]. The study of the progression of AF (identified as a standing open problem [22]) however necessarily requires a greatly expanded time scale that has thus far not been realized. To address this, we couple an established model of atrial electrophysiology [24], with our proposed model of dynamic control of ion channel expression and intercellular coupling based on *Ca*_*i*_ feedback. The new model opens up the study of cardiac activity across multiple timescales, from beat-to-beat dynamics to the slow progression of substrate changes and AF progression, paralleling developments in the long-term study of neuronal homeostasis and plasticity [25], the development of historical structures [26], or more generally of multiple timescale processes in adaptive networks [27].

We find that the single atrial myocyte responds dynamically to increased pacing rates, remodeling ion channel expression and reproducing the characteristic cellular scale changes in APD and CaT morphology well-established in AF. At the tissue scale, while the non-remodeled healthy baseline atrial tissue is resistant to re-entrant electrical activity, continuous rapid pacing leads to tissue remodeling, and thus to the stabilization of re-entrant waves. Critically, we find that fast pacing in a heterogeneous tissue qualitatively replicates the natural history of AF: Pacing induces an initial substrate remodeling, followed by spontaneous re-entry. This paroxysmal AF induces further remodeling, which transforms the intermittent arrhythmic activity into persistent chaotic activity, or permanent AF. Throughout this process, we track the characteristic long-term organization of AF [28], observing an increase in AF episode duration and re-entrant wave locations and their spatial stabilization. Overall, we propose a novel theoretical framework that necessarily ties together multiple interrelated processes (ion channel co-regulation, calcium homeostasis, AF natural history), demonstrating the first in-silico investigation of the entire progression (i.e., initiation to remodeling to stabilization) of AF.

## 2 Methods

### Atrial electrophysiology and calcium feedback regulation model

We simulate individual atrial myocyte electrical dynamics using an established human atrial model of ionic currents and intracellular calcium (*Ca*_*i*_) cycling [24]. To represent long-term electrical remodeling, we couple the atrial electrical and *Ca*_*i*_ model with a *Ca*_*i*_ regulatory feedback model governing ionic current expression, originally proposed for neurons [6] and recently used by us to probe long-term emergent behavior in the sinoatrial node in the heart [7]. Thus, in this coupled system, on the short-term (milliseconds to seconds) timescale, atrial transmembrane potential (*V*) is governed by voltage (*V*)- and *Ca*_*i*_-dependent ionic currents *I*, while calcium currents and fluxes govern *Ca*_*i*_ [24]. On the long-term (minutes to hours / days) timescale, ionic conductances (which mediate ionic currents and fluxes), are governed by the *Ca*_*i*_ feedback model.

In brief, the fundamental mechanism underlying the *Ca*_*i*_ regulatory feedback model is that individual myocytes regulate ion channel expression to ‘match’ an inherent intracellular calcium target level (*Ca*_*tgt*_) to maintain *Ca*_*i*_ homeostasis. The feedback model is formulated as

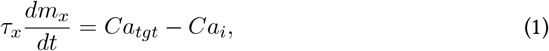

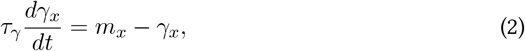

where *Ca*_*tgt*_ is the homeostatic calcium target, *Ca*_*i*_ is the intracellular calcium concentration, *m*_*x*_ represents a relative (normalized) mRNA expression compared to baseline, and *γ*_*x*_ is a scaling factor for ionic conductances *g*_*x*_, for each of the *x* ionic currents *I* and fluxes *J* coupled to the feedback system, which are those with significant changes (i.e. mRNA, protein expression or current density) in AF remodeling (*I*_*Na*_, *I*_*CaL*_, *I*_*to*_, *I*_*Kur*_, *I*_*K*1_, *I*_*Ks*_, *I*_*NCX*_, *J*_*RyR,leak*_, *J*_*SERCA*_) [3]. Notably, time constants *τ*_*x*_ and *τ*_*γ*_ determine the timescale of the response to *Ca*_*i*_ changes.

We simulate a two-dimensional atrial tissue by electrically coupling individual atrial myocytes, represented by the standard monodomain partial differential equation formulation, with diffusion coefficient *D* governing cell-cell electrical coupling. We choose a spatial discretization step of Δ*x* = 0.125 cm, corresponding to the average size of a human atrial myocyte [29]. The largest tissue size we simulate (1024 *×* 1024 cells or 12.8 *×* 12.8 cm) is equivalent to the total area of both atria [30]. To represent long-term intercellular coupling remodeling (reduced gap junctional coupling, fibrosis) in AF, *D* is also scaled by *γ*_*D*_, which similarly obeys equations Eqs. 1-2. Importantly, note that in tissue, each cell has a local *Ca*_*tgt*_ and its own set of *γ*_*x*_ (for all conductances and diffusion), coupled to the local *Ca*_*i*_, such that all *γ*_*x*_ vary in space, in addition to time.

In the healthy state (i.e., absence of remodeling), *γ*_*x*_ = 1 for all of currents, fluxes, and intercellular coupling. While *τ*_*γ*_ is equal for all ionic conductances (approximately representing the timescale of protein translation), crucially, *τ*_*x*_ is unique for each conductance, thus defining their individual dynamic response to changes in *Ca*_*i*_. We derive the *τ*_*x*_ values from the healthy and the chronic AF values based on measured AF electrical remodeling changes (see Table 5 in ref. [3]), as shown in detail in the Supplement. Importantly, noting that intercellular coupling remodeling occurs on an approximately 1000-fold slower timescale than electrical remodeling [31], *τ*_*D*_, the time constant for *γ*_*D*_ in Eq. 1, is three orders of magnitude larger compared with the time constants for ionic currents and fluxes. Further, we choose the baseline *Ca*_*tgt*_ of 258 nM, to be equivalent to the time-average *Ca*_*i*_ in the atrial cell model paced at 60 bpm (1000 ms cycle length), thus defining the baseline homeostatic state. In heterogeneous tissue simulations, we vary *Ca*_*tgt*_ on an interval [200 − 320] nM, assuming approximately 20% tissue heterogeneity in the healthy atria.

Finally, we note that simulating the long-term changes associated with AF remodeling poses a significant computational challenge. To solve this system efficiently, we developed a CUDA-based model implementation to utilize state-of-the-art GPU hardware available through the Ohio Supercomputer Center [32] and the Delta cluster at the National Center for Supercomputing Applications, through an NSF Access allocation [33]. The GPU formulation takes advantage of the massively parallel nature of the system and enabled investigation of continuous electrical activity in a large tissue for more than 24 hours of simulated time - to our knowledge, the longest cardiac electrophysiology simulations performed thus far.

A complete description of the model formulation, parameter derivation, tissue pacing protocols, spiral wave tracking, action potential (AP) characterization, heterogeneous *Ca*_*tgt*_ maps, and numerical methods is provided in the Supplement.

## 3 Results

### AF remodeling following rapid pacing in a single atrial myocyte

We first investigate the behavior of a single atrial cell coupled to the feedback model. Individual cell ion channel conductances remodel in response to long-term pacing at different cycle lengths (Figure 1A). At a cycle length of 1000 ms (60 bpm), the average resting heart rate in humans, all conductance scaling factors (*γ*_*x*_) remain close to their nominal value of 1, consistent with homeostasis (i.e., *Ca*_*tgt*_ = 258 nM) corresponding to baseline channel expression. In response to faster pacing, i.e., shorter cycle length, *m*_*x*_ is modified (see Figure S1), driving changes in conductances *γ*_*x*_, following Eqs. 1-2, which thus dynamically adapt and reach a new steady state. Note that some conductances increase (*γ*_*NCX*_, *γ*_*RyR*_*leak*__, *γ*_*Ks*_, *γ*_*K*1_) while others decrease (*γ*_*Na*_, *γ*_*SERCA*_, *γ*_*ito*_, *γ*_*CaL*_, *γ*_*Kur*_), based on their respective up- or down-regulation during AF remodeling (described further in the Supplemental Methods). Importantly, the long-term conductance dynamics are driven by *Ca*_*i*_ (Figure 1B): faster pacing initially increases the *Ca*_*i*_ average (solid black curve) above the cell’s inherent *Ca*_*tgt*_ (dashed black line), thus driving the remodeling (i.e., change in conductance scaling factors) via the feedback model. This subsequent remodeling ultimately leads to a decrease in the average *Ca*_*i*_, until the time average reaches *Ca*_*tgt*_, thus maintaining homeostasis. Progressively faster pacing leads to progressively greater remodeling; notably, the steady-state following pacing at the fastest cycle length of 250 ms leads to conductances near levels in chronic AF (see Table S1). The remodeling process is also reversible: the scaling factors return to their original value of 1 following a return to pacing at 1000 ms.

**Figure 1.**
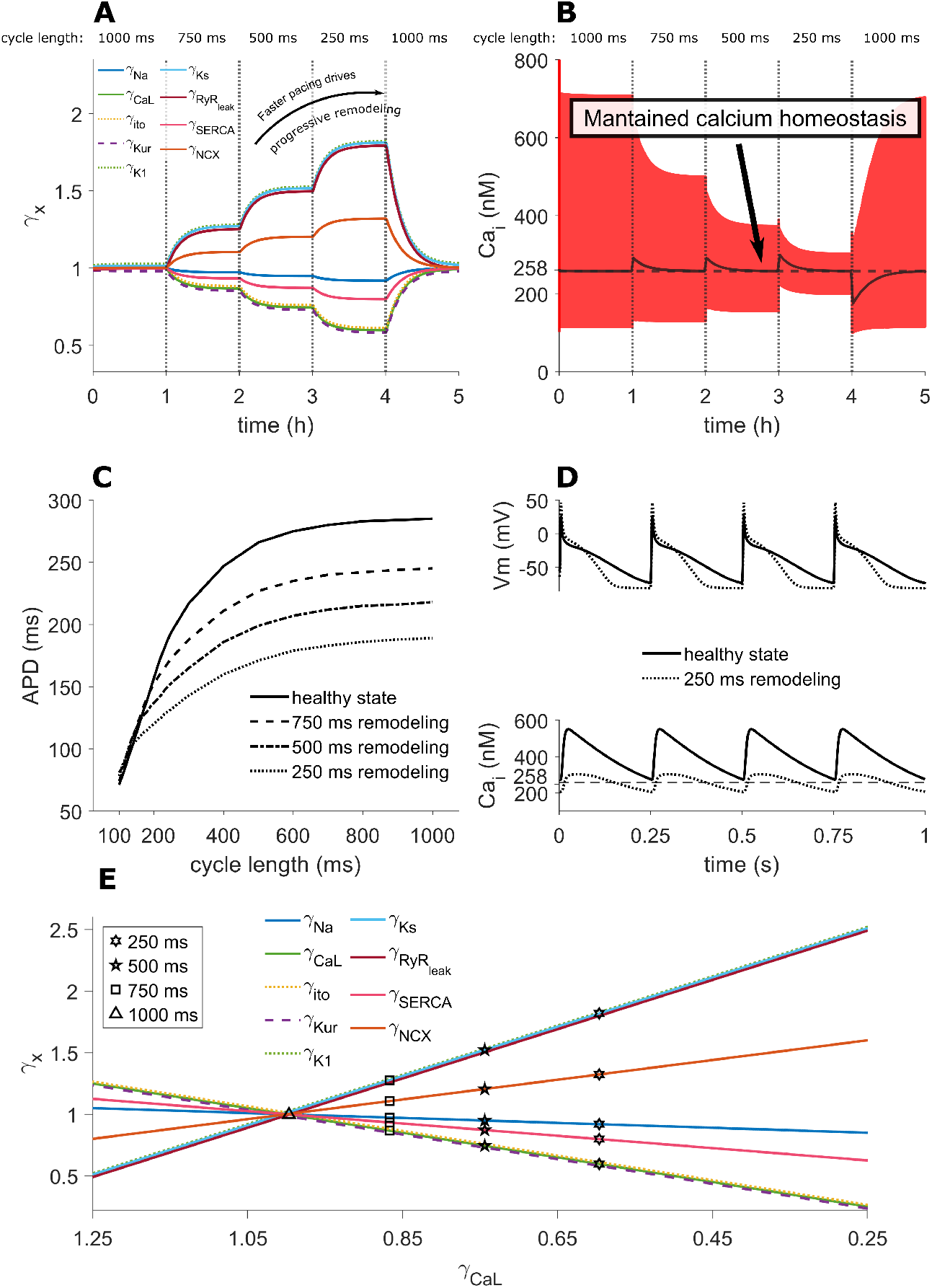
Electrical remodeling in individual atrial myocytes. **A**. Progressive electrical remodeling of ionic currents and fluxes (different color lines) in response to progressively faster pacing (shorter cycle lengths). Conductances are at baseline (*γ*_*x*_ *≈* 1) for pacing at 60 beats per minute (cycle length of 1000 ms). **B**. Calcium transients (CaTs, red) and moving average *Ca*_*i*_ (black) illustrate the impact of remodeling to a maintain a homeostatic *Ca*_*i*_ level (*Ca*_*tgt*_, dashed horizontal line) at different cycle lengths **C**. Electrical remodeling, following pacing at different cycle lengths, alters APD restitution. **D**. Electrical remodeling following 250 ms pacing (dotted line) results in shorter action potentials (AP) and altered CaTs, compared with the healthy state (solid line) (*Ca*_*tgt*_, dashed horizontal line). **E**. For all currents and fluxes (different color lines9), steady state remodeling at different cycle lengths (symbols) maintains a linear relationship between conductances, described by Eq. 1.

We measure the characteristics of action potentials (APs) and CaTs following progressive remodeling (i.e., the steady-state for progressively faster cycle lengths). Notably, progressive remodeling leads to the characteristic electrical changes found in chronic AF: a flattening of the APD restitution curve (Figure 1C), shortened APD, lower resting membrane potential, and reduced CaT amplitude (Figure 1D). Reduced CaT are a crucial result of the feedback system: faster pacing intrinsically elevates *Ca*_*i*_, such that remodeling necessarily reduces the CaTs to maintain long term *Ca*_*i*_ homeostasis towards *Ca*_*tgt*_ (dashed black line). Additionally, while the overall dynamics are similar, the specific *Ca*_*tgt*_ level alters the steady state conductances at the different cycle lengths (Figures S2-S4), with lower *Ca*_*tgt*_ leading to more remodeling and vice-versa. Notably, higher *Ca*_*tgt*_ results in a steeper APD restitution curve in response to remodeling at all long term pacing rates (Figure S4).

Figure 1E illustrates channel conductances relative to the reference *γ*_*CaL*_, with the markers denoting the steady state values, highlighting their respective linear relation maintained throughout remodeling. Notably, this illustrates that for any *γ*_*CaL*_ value, there is a corresponding set of all other conductances, uniquely defined by the linear relationship. Therefore, for simplicity, in the subsequent tissue simulations, we present a single scaling factor (*γ*_*CaL*_), recognizing that all other conductances will be either proportional (if similarly down-regulated as *γ*_*CaL*_) or inversely proportional (if up-regulated). We note the relative relationship between the conductances is determined by the *τ*_*x*_ values derived from experimental data, as explained in Supplemental Methods. Further, it is notable that while the *τ*_*x*_ values define the direction and relative magnitude of conductance changes, they do not impose the steady-state at a given cycle length. Therefore, it is an emergent property that the steady-state following rapid pacing of 250 ms is remarkably close to the remodeling observed in chronic AF.

Additionally, we demonstrate that the model is robust to both variation in the *τ*_*x*_ values (Figure S5), where random *τ*_*x*_ values correspond to random slopes for the linear relationships shown in Figure 1E, and in the *γ*_*x*_ initial conditions (Figure S6). Importantly, the feedback model enables adequate response to fast pacing, inducing steady-state AP and CaT morphology that are generally consistent with AF remodeling. Overall, the single cell model captures key dynamics of electrical remodeling in AF: the ion conductance responses to rapid pacing, subsequent changes in AP and CaT properties, and reversibility.

### Progressive AF remodeling in homogeneous atrial tissue

We next investigate the behavior of re-entrant activity in a two-dimensional atrial tissue, initially considering a homogeneous tissue (Figures 2, S8), in which all cells have identical *Ca*_*tgt*_ = 258 nM, as in the single cell simulations above. We apply a protocol designed to investigate the effect of long-term re-entrant activity in tissue and specifically the interaction between that activity and remodeling: The tissue is first rapidly paced (10 seconds at 100 ms cycle length) to drive remodeling. Then, we induce a spiral wave by cross-stimulation and track the persistence of re-entrant activity over time. If electrical activity is extinguished (i.e., the spiral wave self-terminates), we restart pacing followed by cross-field stimulation (see Supplemental Methods for complete protocol details).

**Figure 2.**
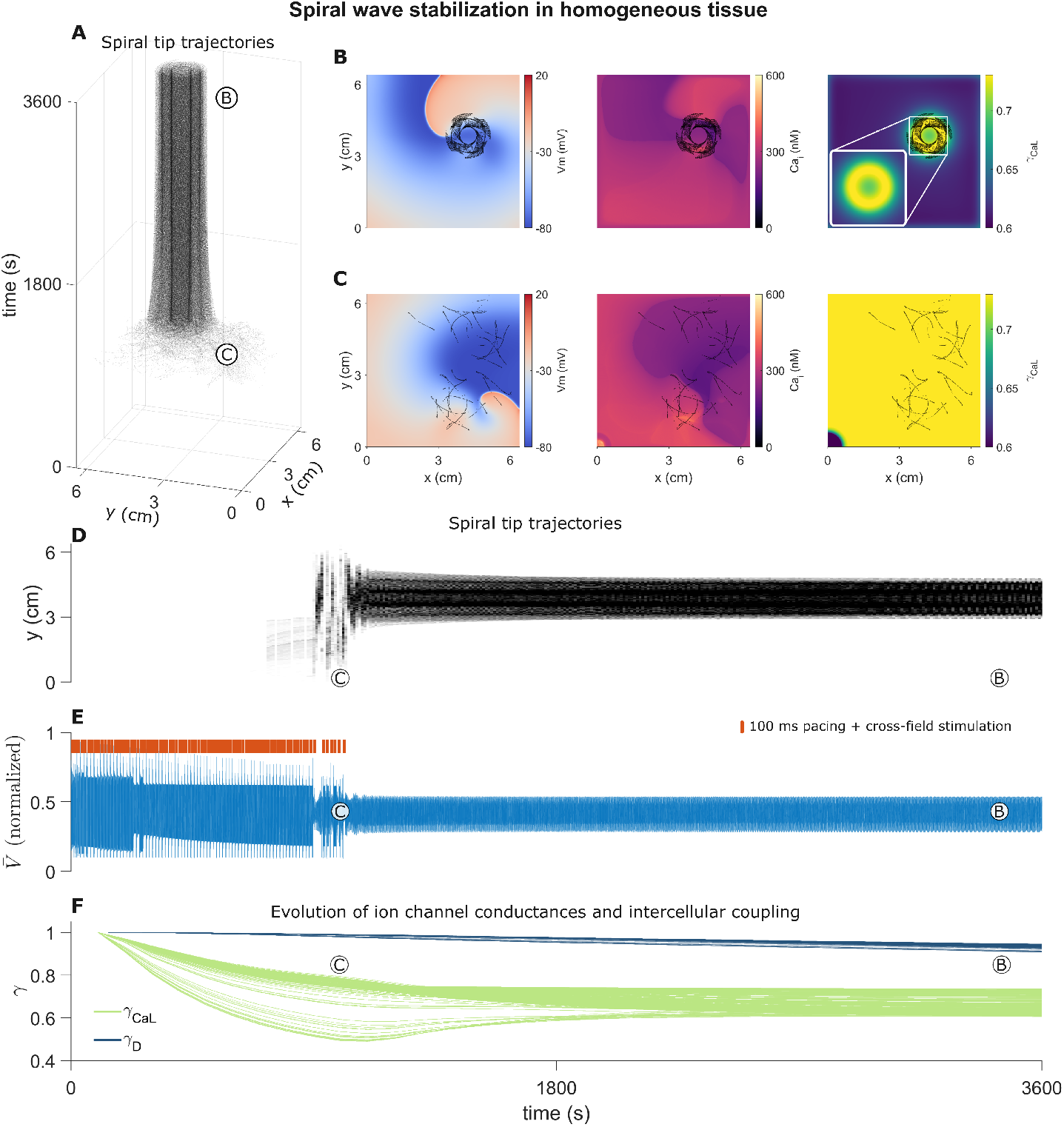
Spiral wave stabilization in a homogeneous atrial tissue. The size of the tissue corresponds to 512*×*512 cells. **A**. The trajectory of the spiral tip over the first simulation hour illustrates stabilization after the first approximately 1000 s. **B, C**. Snapshots of voltage, *Ca*_*i*_ and electrical conductance scaling factor *γ*_*CaL*_ for (B) stable and (C) unstable spirals. The initially induced spiral wave is unstable, leading to breakups and extinction. As remodeling progresses, the spiral wave stabilizes. Note the spiral tip follows the electrical remodeling spatial pattern, which was induced by the stable spiral wave (B, right panel and inset). Black traces show spiral tip position shortly before and after each snapshot. **D**. The position of the spiral tip trajectories (y-axis projection) is initially wide (spanning the entire tissue), with brief pauses (denoting wave breakups), before stabilizing. **E**. The pseudo-ECG of tissue voltage activity similarly indicates periods of unstable and stable spirals. Orange lines denote rapid (100 ms) pacing followed by cross-stimulation. **F**. Electrical (*γ*_*CaL*_, green lines) remodeling approaches steady-state levels relatively early, compared with the slower intercellular coupling (*γ*_*D*_, blue lines) remodeling progression on a longer timescale. Lines denote *γ* _*CaL*_ and *γ* _*D*_ at each spatial location in the atrial tissue. Note that, as explained in text, *γ*_*CaL*_ represents all dynamic conductances.

The spiral tip trajectory (i.e., the position of spiral tip(s) over time) illustrates that during the first hour of simulation, electrical activity transitions from an unstable pattern, with chaotic rotor tip trajectories and multiple spiral wave breakups, to a single stable spiral wave (Figure 2A). Snapshots of tissue voltage at the end and the start of the first hour (Figure 2B and C, left, respectively) illustrate the stable wave pattern that emerges. In addition to the chaotic trajectories, the initially induced re-entrant activity frequently self-terminates, leading to a restart of rapid pacing (see gaps in trajectories in Figure 2D, orange lines - denoting pacing episodes - above the tissue pseudo-ECG in Figure 2E).

Importantly, spiral wave stabilization is driven primarily by the electrical remodeling (Figure 2B, C, right; F, green lines). The initial episodes of fast pacing and the transitory spiral waves induce sufficient tissue remodeling, developing a substrate that can sustain stable activity. Beyond the tipping point for this substrate, a single stable spiral wave forms. The stabilized spiral wave then further rapidly paces the tissue, sustaining the remodeling process until a new steady state is reached.

A surprising emergent result (and prediction) of this tissue simulation is that remodeling is spatially heterogeneous, despite identical cells within the tissue having a single homogeneous *Ca*_*tgt*_ (see Figure 2B, right, inset). Specifically, the region in the atrial tissue corresponding to the location of the stabilized spiral wave tip trajectory is the *least* remodeled (i.e., electrical conductances closest to 1), with a ‘ring’ spatial pattern with reduced remodeling emerging. The formation of this remodeled ‘niche’ is caused by the fact that cells at and near the spiral wave ‘core’ are effectively only partially activated, compared to the rest of the tissue, which induces less remodeling. Also notably, on this timescale, there is minimal remodeling of intercellular coupling, i.e., *γ*_*D*_ remains near 1 throughout the tissue (Figure 2F, blue lines), demonstrating that this niche formation is driven initially by electrical remodeling.

Simulating beyond the first hour, we find that the location of the spiral wave initially remains consistent (Figure S8A, see also Movie S1). However, subsequent intercellular remodeling induces a long term, slow drift of the spiral wave core (Figure S8A). Notably, this pattern is different from other forms of spiral meandering, as the immediate pattern of re-entry remains quasi-stable, and only drifts on a very slow time scale. The ring remodeling spatial pattern is initially apparent in the *γ*_*D*_ map as well, but the pattern becomes ‘blurred’ by the re-entrant wave drift (Figure S8D, E, two right panels). Further, the overall level of electrical remodeling is maintained (Figure S8C, green lines), and the intercellular remodeling also reaches a remodeled state during the 24 hours (Figure S8C, blue lines). Notably, the changes in the underlying substrate have a complex effect on the spiral trajectory itself, with a transition to a different pattern in the last quarter of the simulation, as evidenced in the spiral tip location y-position projection in panel A, as well as the traces in panels D and E.

Different *Ca*_*tgt*_ levels within the homogeneous tissue have a significant influence on long-term tissue level re-entrant behavior. Using the same pacing and cross stimulation protocol, a lower *Ca*_*tgt*_ of 200 nM (Figure S9, Movie S2) leads to more remodeling (lower *γ*_*CaL*_, *γ*_*D*_), consistent with single cell behavior (Figure S2). Enhanced remodeling shortens APDs in the tissue and results in significantly more dynamic, chaotic behavior. Following a shorter period of pacing, two spiral waves are induced. As the remodeling progresses, the spirals change location in the tissue, ultimately leading to a brief pause in re-entrant activity (Figure S9B, close to the 16 hour mark). Pacing and cross-field stimulation then induces a single spiral wave which stabilizes on the previously remodeled ring pattern of an extinguished re-entrant wave (Figure S9F, Movie S2).

Conversely, the same protocol applied to a homogeneous tissue with higher *Ca*_*tgt*_ of 320 nM leads to less remodeling (Figure S10, Movie S3), also consistent with single cell behavior (Figure S3). This in turn leads to the inability of cross-wave stimulation to induce stable re-entry. Every induced wave is unstable and quickly extinguishes, leading to a restart of the pacing protocol.

To separate the effects of electrical vs. intercellular coupling remodeling, we simulate a homogeneous tissue with the baseline *Ca*_*tgt*_ = 258 nM while ‘clamping’ (i.e., fixing as a constant) *γ*_*D*_ to its initial baseline value of 1 (Figure S11, Movie S4). The overall long-term behavior is comparable to the early activity in the tissue with unclamped *γ*_*D*_ dynamics (Figure S8): initial intermittent re-entry followed by pacing, which then stabilizes as a persistent spiral wave. Notably, the spiral wave remains stable, without either long-term drift or breakup. It is interesting to note that *γ*_*CaL*_ approaches steady state values before re-entry becomes persistent. These results underlie a key model prediction: electrical remodeling is the initial stabilizing factor for re-entry, while the slow intracellular remodeling drives the transition into more complex drifting or chaotic behavior.

### AF progression in heterogeneous atrial tissue

Finally, we investigate the progression of AF in a large, heterogeneous tissue (Figures 3 - 6, Movie S7), such that the average *Ca*_*tgt*_ value within the entire tissue is the same as the homogeneous tissue above (258 nM), while the *Ca*_*tgt*_ value at each spatial location (i.e., each atrial cell) has a different value on the interval [200, 320] nM (Figure 3 E). The tissue is similarly paced at the same 100 ms cycle length (Figure 3 F), but without externally inducing the spiral wave through cross-stimulation. Following pacing for a short period (*≈* 650 s), sufficient remodeling is induced, such that a spiral wave spontaneously forms (Figure 4, Movie S5): spatial heterogeneity in *Ca*_*tgt*_ leads to heterogeneous remodeling and thus to local conduction block, around which re-entry forms. Initially, re-entry is unstable, leading to frequent spiral wave extinction and subsequent restart of pacing (Figure 3 A, B).

**Figure 3.**
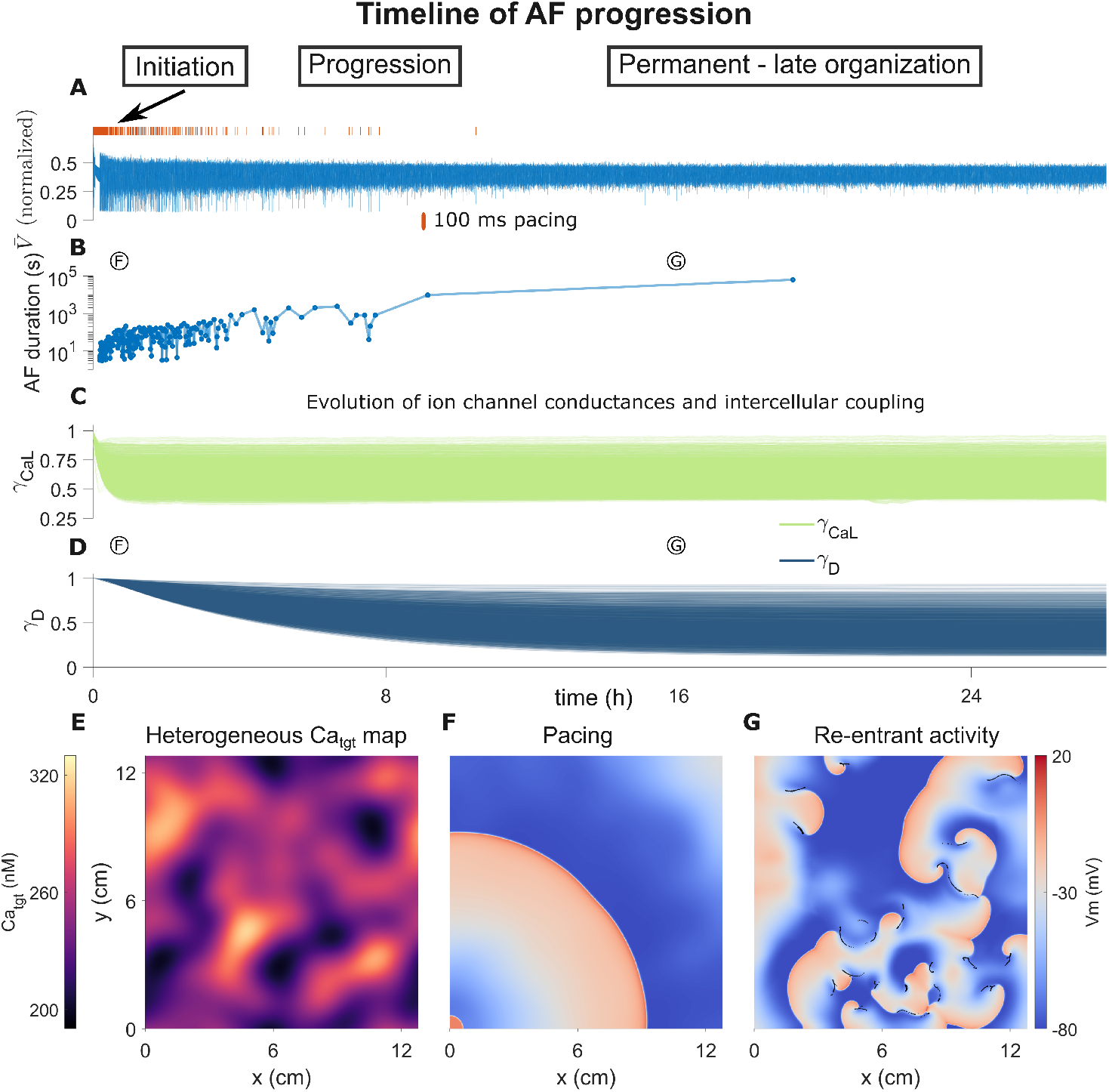
Long term progression of atrial fibrillation in a heterogeneous atrial tissue. Further details of the same simulation are shown in Figures 4 - 6. The tissue size corresponds to 1024*×*1024 cells. **A**. Pseudo-ECG of tissue voltage activity. Orange lines mark episodes of fast pacing (100 ms). **B**. Progression of AF interval duration. Note the log scale on the y-axis. **C**. Electrical (*γ*_*CaL*_) and **D**. intercellular coupling (*γ*_*D*_) remodeling dynamics during progression from transient to persistent fibrillatory activity. **E**. Spatial map of *Ca*_*tgt*_ in the heterogeneous tissue. **F, G**. Examples of electrical activity (time points denoted in the plots above) during an episode of fast pacing and during re-entrant activity.

**Figure 4.**
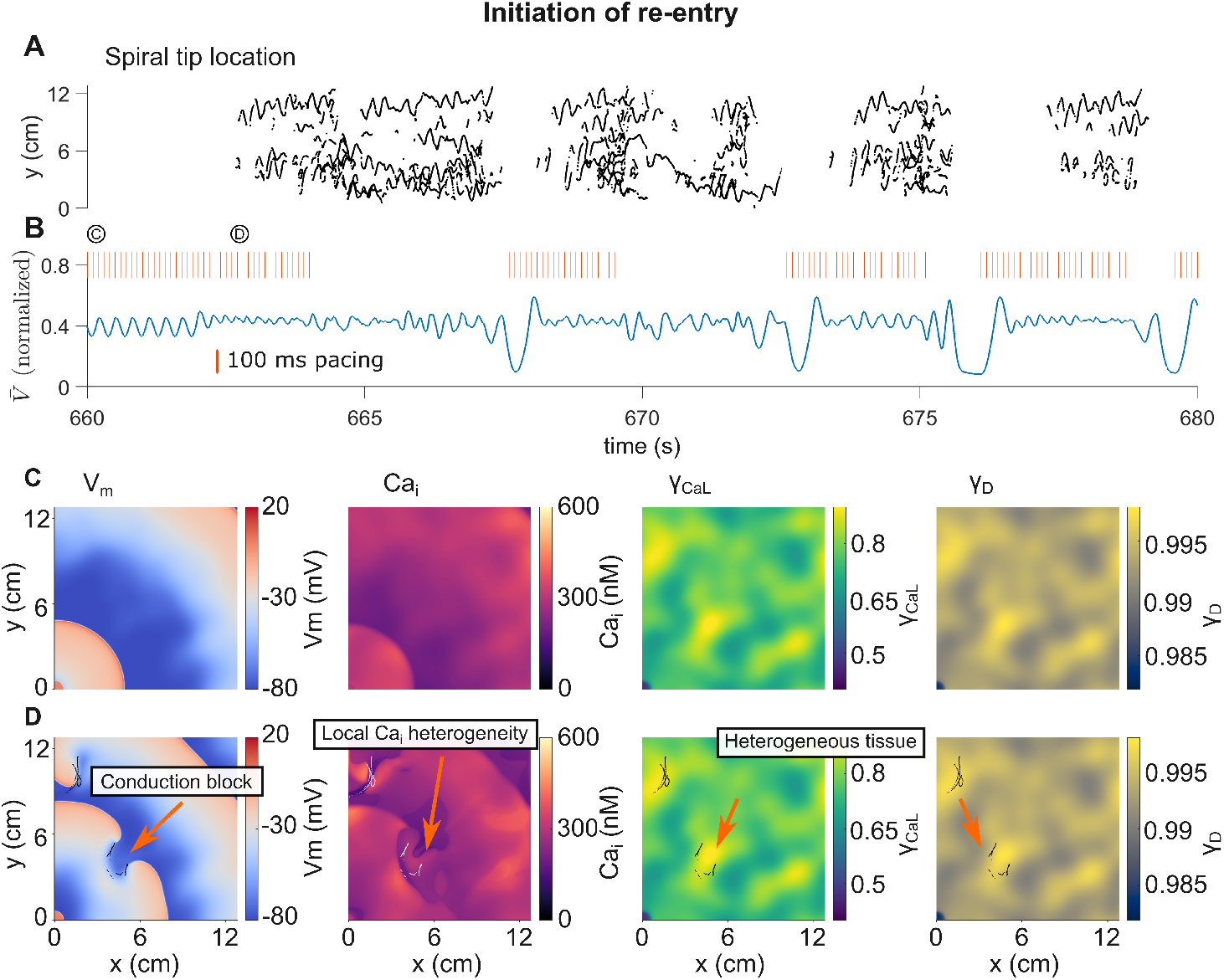
The first episode of spontaneous initiation of re-entry in heterogeneous tissue. **A**. Spiral tip tracks during the first episode of re-entry. **B**. Pseudo-ECG and pacing episodes. **C, D**. Tissue activity and remodeling states during the initiation of re-entry. Panel C illustrates pacing in the lower left corner. Panel D shows re-entry initiation around a remodeling ‘peak’, which defines a local *Ca*_*i*_ heterogeneity leading to a temporary local conduction block. Data is also shown in Movie S5

Electrical remodeling in the atrial substrate drives the initial capability for re-entry initiation (Figure 4D), as the early changes in intercellular coupling are minimal (Figure 4E). However, the pacing and the subsequent re-entrant episodes also drive changes in *γ*_*D*_ (Figure 3D), occurring in parallel with both increasingly longer re-entrant episodes (Figure 3B) and an increasing number of spiral waves that can be sustained in the tissue (Figure 5A). The progression of tissue remodeling further leads to an increase in the dominant frequency of tissue voltage activation, as well as to faster and more complex activity (Figure 5B, C). Ultimately, re-entrant activity becomes permanent. The re-entrant waves in turn maintain *γ*_*CaL*_ and *γ*_*D*_ at a remodeled steady state, capable of sustaining permanent arrhythmic activity.

**Figure 5.**
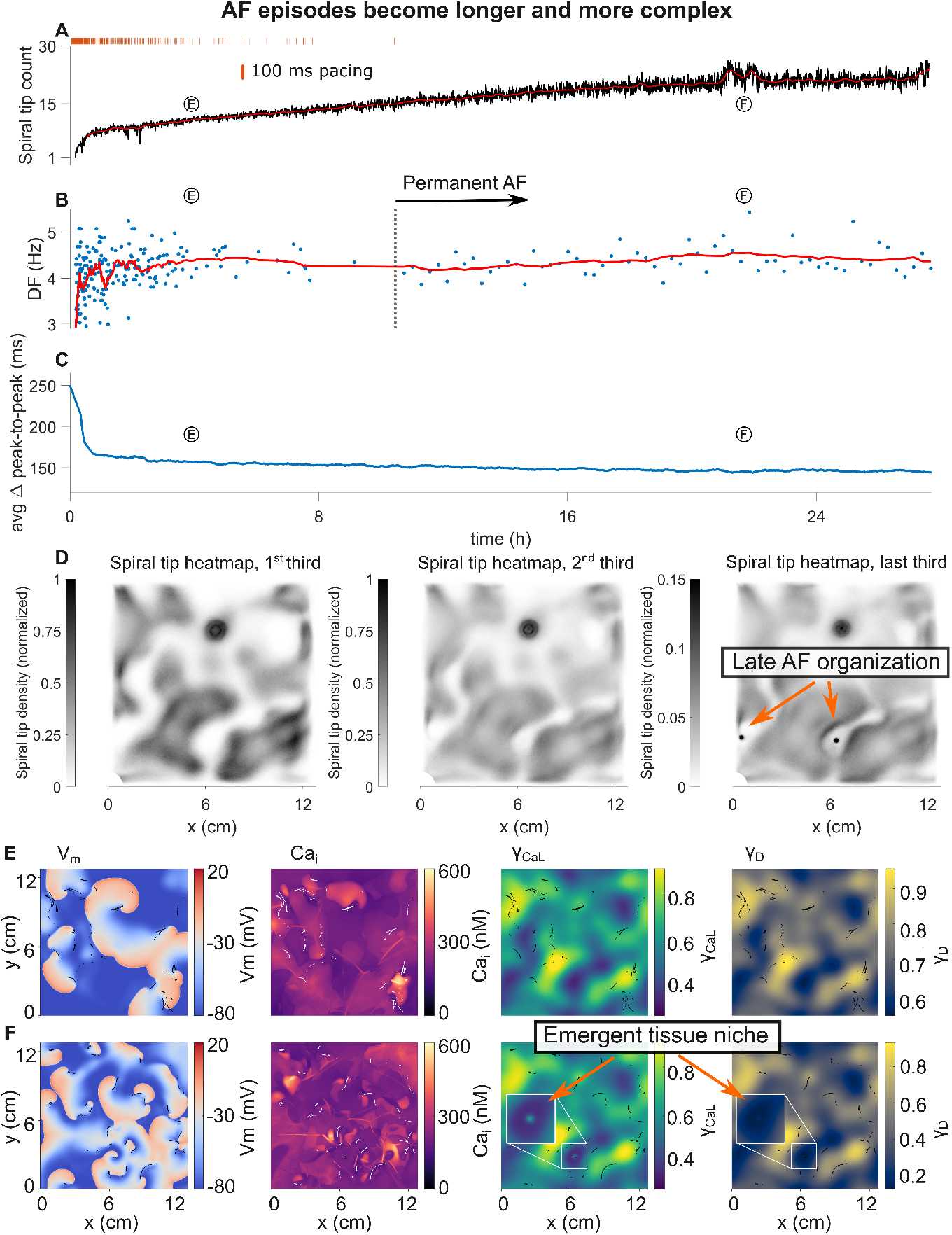
Increasing complexity and organization of AF over time. **A**. The number of spiral wave tips present in the atrial tissue (with red trace showing moving average). The pacing markers (same as Figure 3 A) are overlaid for reference. **B**. Increase in the dominant frequency of AF activity. The vertical dotted line marks the transition to permanent reentrant activity. The final permanent episode is split into 1000 s intervals. The red curve represents the moving average of the frequency, increasing over time. Note that the y-axis was chose to clearly delineate the average, such that some data points are outside the plot. **C**. Moving average of distance between peaks in the pseudo-ECG, a measure of fractionation or complexity of activation. **D**Spiral tip density heatmap of each third of the simulation. **E**,**F**. Snapshots of voltage, *Ca*_*i*_, *γ*_*CaL*_, and *γ*_*D*_ at time periods corresponding to (E) early and (F) late chaotic electrical dynamics. Black and white lines show spiral tip position shortly before and after each snapshot. Insets in F illustrate the remodeling pattern underlying the stable spiral location.

Notably, the spatial pattern of spiral wave tips is dynamic, as demonstrated in the heatmaps of spiral wave tip distribution during each third of the simulation in Figure 5D. At both early and late time points within AF progression, the spiral tip density is greatest around the locations of transitions between high and low remodeling (Figure 5E, F, two right panels for spatial maps of *γ*_*CaL*_ and *γ*_*D*_). As remodeling progresses, spiral waves are present over a larger amount of the tissue. The spiral tip density exhibits exhibits an interesting spatial pattern during the last third of the simulation (Figure 5D, leftmost panel): stable re-entrant spirals appear in the low *Ca*_*tgt*_ areas, with a small reentry core similar to the simulations in homogeneous tissue (Figure S9). This pattern is accompanied by a corresponding positive ‘bump’ in remodeling underlying the wave location (Figure 5F, inset), reminiscent of the pattern that develops in homogeneous tissue. Due to the fast activation of this re-entrant spot, the density of other spiral waves in the nearby transition zone increases (Figure 5D), and further lead to an increase in total number of re-entrant waves the tissue can sustain (Figure 5A). This pattern is not completely stable, and appears in two low *Ca*_*tgt*_ zones, as highlighted in Movie S7.

We also investigate the state of the fully remodeled atrial tissue, at the end of the simulation (Figure 6, Movie S6). An emerging stable re-entry site is apparent in the location of the spiral tips (Figure 6A, orange line). This leads to the emergence of a niche as described above - note the positive ‘bump’ in the *γ*_*CaL*_ inset in Figure 6C. This fast localized activity is evident as a small, bright spot in the tissue activation frequency map (Figure 6D, white arrow). The area immediately close to the stable re-entrant focus exhibits the most frequent activation. Notably, throughout the tissue, the electrical activity exhibits multiple frequencies and is highly dissociated from calcium activity, hallmarks of AF. In all cells shown in Figure 6, calcium activity settles in a complex alternating pattern, with large peaks followed by multiple smaller transients, with each spatial location ultimately maintaining their respective *Ca*_*tgt*_.

**Figure 6.**
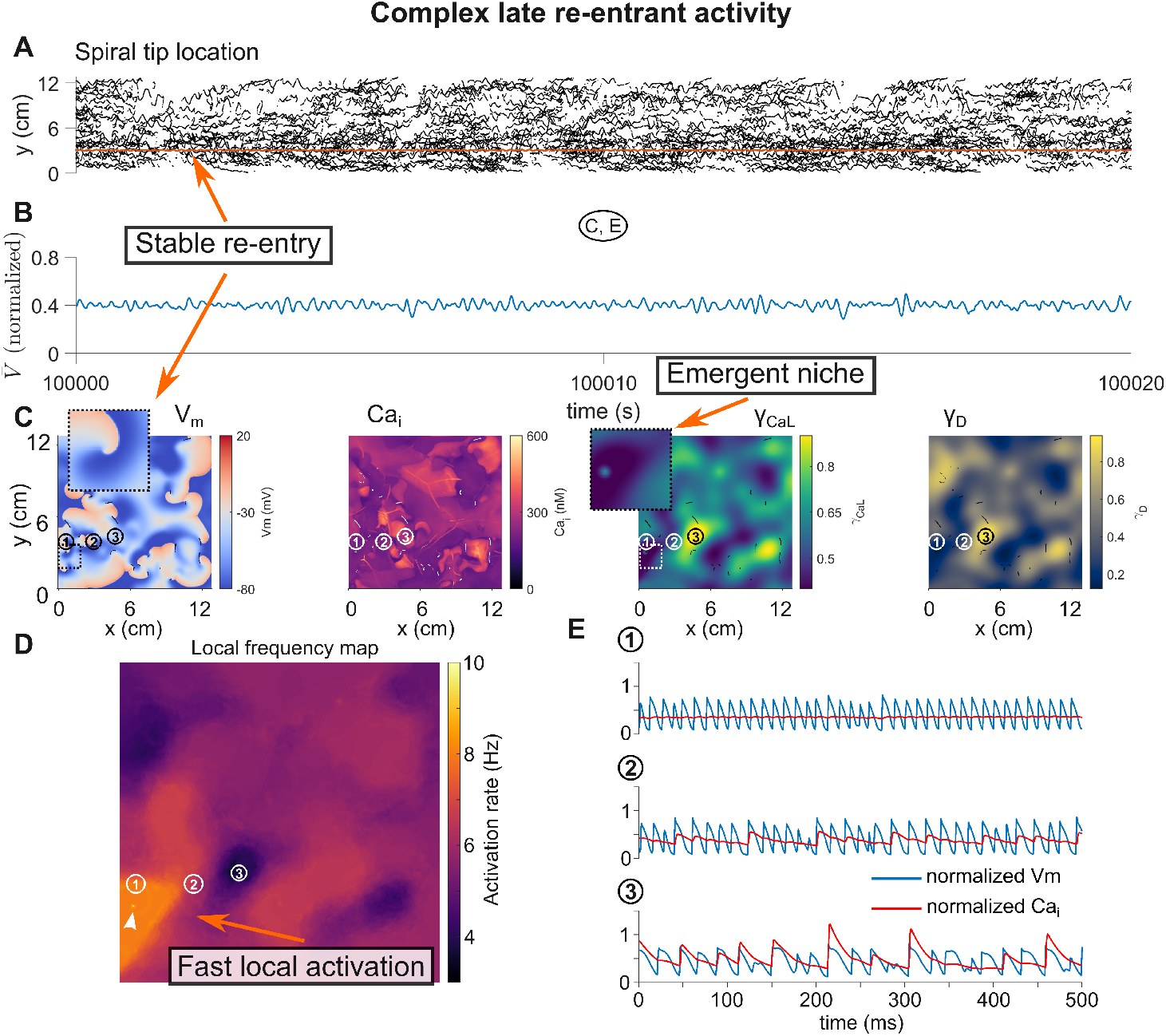
Tissue activity in fibrillatory tissue. **A**. Dense spiral tracks during fibrillatory activity. The orange track highlights the stable re-entrant activity in the emergent tissue niche. **B**. Chaotic tissue pseudo-ECG. **C**. Snapshot of tissue activity and remodeling state. Points 1, 2 and 3 mark the cell locations highlighted in panel E. White arrow denotes the stable spiral location. **D**. Map of average voltage activation frequency, calculated from the 20 s of activity shown in panels A, B. White arrow points to high spot created by stable spiral wave location, which rapidly paces the local region. **E, 1, 2, 3**. Normalized voltage (blue) and calcium (red) activity at the three locations. Cell 1 exhibits the most remodeling, with the highest activation frequency and very low calcium activity. Cells 2 and 3 are less remodeled, showing high calcium peaks and longer APDs. Both are activated chaotically and exhibit APD and calcium peak irregular periodicities. Note the dissociation between voltage and calcium, a hallmark characteristic of atrial fibrillation (AF).

Finally, we compare the fully AF remodeled tissue with the corresponding “control” tissue (i.e., same *Ca*_*tgt*_ map) that has instead been paced at 1000 ms (60 bpm) (Figure S12), uncovering significant changes in tissue properties. Throughout the tissue, APD values in the AF remodeled tissue are significantly shorter (Figure S13A, B), and with a wider normalized distribution (Figure S13 E). Conduction is also slower and more irregular in the AF tissue (Figure S13C, D). Ultimately, these changes lead to an increased susceptibility of AF tissue to re-entry (Figure S14). Notably, while tissue re-entry can be induced in the control tissue, the episodes are brief and self-terminating; in contrast, in AF remodeled tissue, a single pacing episode induces permanent re-entry.

Importantly, the activity described in Figures 3 - 6 is only one out of a number of possible electrical activity and remodeling spatiotemporal patterns: We tested the same pacing protocol on 25 different random *Ca*_*tgt*_ maps (Figures S15-S16), each with the same range of *Ca*_*tgt*_ values but with a different spatial pattern. Note the case described above is number 8 out of the 25 (with arbitrary ordering). We find that 3 patterns of activity emerge: (i) no re-entrant activity is induced, and therefore pacing is continuous (6 cases), (ii) transitory re-entry followed by pacing (13 cases), and finally (iii) persistent re-entrant activity (6 cases). An example of a tissue resistant to re-entry and intermittent re-entry are shown in Movies S9, S10, corresponding to maps 2 and 3 respectively, and two further examples of persistent activity are shown in Movies S8, S11, corresponding to maps 1 and 5. Thus, a majority of tissues present with either transient or persistent re-entrant activity, with the exact spatiotemporal pattern determined by the heterogeneous *Ca*_*tgt*_ map. Notably, in the simulations with permanent re-entry shown (Movies S8, S11), the same distinct spiral density hotspots appear in low *Ca*_*tgt*_ areas, towards the end of the simulation, which also induce a remodeling ‘bump’ pattern as described above.

## 4 Discussion

### Model assumptions

We first discuss several critical model assumptions. Figure 1 illustrates the fundamental assumptions and dynamics of the regulatory model, which are imposed at the level of the single atrial cell. First, the regulatory model maintains constant average *Ca*_*i*_ levels, determined by the *Ca*_*tgt*_ set point, by dynamically adjusting mRNA and thus ion channel expression. As discussed in our previous work [7], the single *Ca*_*tgt*_ is a reduction of a highly complex intracellular regulatory network [5] to a single phenomenological setpoint. There are multiple known mechanisms through which pacing-induced increases in *Ca*_*i*_ translate to ion channel changes: increased *Ca*_*i*_ activates calcineurin, a Ca^2+^-dependent phosphatase, which activates nuclear factor of activated T cells (NFAT) that suppresses Cav1.2 (channel carrying *I*_*CaL*_) transcription [34]. Moreover, NFAT upregulates *I*_*K*1_ mRNA and protein expression by removing translational inhibition [35] and further reduces *I*_*to*_ mRNA and protein expression [36]. Increased *Ca*_*i*_ additionally leads to activation of CaMKII, which can decrease sarcoplasmic reticulum Ca^2+^ uptake and increases ryanodine receptor leak [37]. Further, the knockout of transcription factor PITX2 leads to AF-like remodeling in the absence of fast pacing [38], suggesting that it could play a part in setting the sensitivity of the regulatory network to *Ca*_*i*_. Taken together, despite their complexity, all these pathways act in concert to reduce average *Ca*_*i*_, similar to the results shown in Figure 1 B. Thus, the model captures the fundamental behavior of the regulatory network - adjusting ion channel expression to maintain calcium homeostasis.

Other pathways could be targeted by intracellular regulation during AF progression. For example, it has recently been shown that Ca^2+^ buffers are significantly decreased in chronic AF [39], leading to an increased risk of spontaneous Ca^2+^ release or arrhythmia initiation. By implementing a similar regulation as utilized to represent ion channel expression, we can reproduce the regulation of Ca^2+^ buffer concentrations in a single cell (Figure S7 I, A). However, we find that incorporating this additional regulation results in minimal changes in subsequent ion channel remodeling (Figure S7 I, B) and AP shape (Figure S7 I, B, top panel), with small differences in the CaT shape (Figure S7 I, B, bottom panel).

In addition to the overall *Ca*_*i*_ level, the balance between diastolic and systolic *Ca*_*i*_ levels is critically important for cardiac contractile function [40], especially given the fast pacing rates that occur in AF. As such, the feedback model can be naturally extended to investigate the influence of regulation during these two phases, with two *Ca*_*tgt*_ values that alternate in time with the phases of the CaT based on average systolic and diastolic *Ca*_*i*_ levels (Figure S7, II A). Note that all channels follow the two alternating *Ca*_*tgt*_ values - imposing two targets to different subsets of the channels will lead to an unstable regulatory system [6]. Interestingly, the two targets lead to less remodeling at fast pacing (Figure S7, II B), which drives significant changes in AP shape compared to the baseline model as well (Figure S7, II C, top panel). Further, the resultant CaT fails to perfectly match either of the two targets and instead settles into an intermediate pattern between the two (Figure S7, II C, bottom panel). Collectively, these examples serve to illustrate that the feedback model can be naturally extended to account for more complex intracellular regulation. Furthermore, as discussed below, a model that includes subcellular detail might be necessary to capture the full range of calcium cycling dynamics.

A second important assumption governing single cell dynamics is that as expression changes *during remodeling*, correlations between channels are maintained (Figure 1E), based on the concept of co-expression and co-regulation of ion channel mRNA and protein levels [8–11]. However, the direction of the expression changes in AF remodeling at the atrial cellular level can differ from those in healthy and ventricular cells. For example, *I*_*Kr*_ (carried by hERG1 protein) remains unchanged in AF [3], despite the lower levels of *I*_*CaL*_ or *I*_*Na*_, with which it is co-expressed [11]. Further, more remodeled states (e.g., the steady state at 250 ms pacing rate) show increased levels of *I*_*Ks*_ and *I*_*K*1_, while *I*_*CaL*_ is decreased, which reverses the relation shown in healthy ventricular cells described by Banyasz et al [12]. Therefore, this suggests that the specific regulatory pathways involved in AF-induced atrial remodeling are different than those setting the baseline healthy state of ventricular cells, with differences arising potentially in both a disease- and chamber-specific manner. However, it is plausible that similar regulatory mechanism governing calcium homeostasis (i.e., altered transcription, co-expression, co-transcription) are involved, thus leading to correlated ion channel expression levels, albeit with different relationships between currents adapted specifically to reducing *Ca*_*i*_ load.

The dynamics of tissue remodeling, shown in Figure S8C and Figure 3C, D, highlight the three distinct timescales in our model: The fastest timescale, on the order of milliseconds, is determined by the beat-to-beat dynamics governing APs and CaTs. The intermediate timescale is determined by electrical remodeling (i.e. the dynamics of *γ*, Figure 3C), set by *τ*_*γ*_ (see Supplementary Methods) and constrained by electrical remodeling occurring significantly slower than the beat-to-beat dynamics. The value of the constant *τ*_*γ*_ is defined such that the remodeled *γ* values reaching a new steady state in approximately one hour (e.g. Figure 2F), based on the fastest described changes in electrical properties (e.g. effective refractory period) [41]. Notably, reported timescales vary: other studies show slower changes in these properties, on the order of days [2, 42]. The slowest timescale governs the intercellular coupling or structural changes (highlighted in Figure 3D), known to occur at an even slower rate than electrical remodeling [43, 44]. Here, we assume that these changes stabilize over a period of 24 hours, a faster response than that seen in the studies cited above. However, as shown previously (by us [7] and Jaeger and Tveito [45]), the separation of timescales is sufficient to capture qualitative behavior in cardiac systems with dynamic substrates. Thus, our timeline of electrical properties and intercellular coupling dynamics, while erring towards the fastest timeline of reported data, is adequate to describe the long term behavior of cardiac tissue remodeling. Thus, the separation of the three timescales enables approximately 24 hours of simulation time to qualitatively reproduce the processes of experimental rapid pacing and AF remodeling, which occur over weeks or months. Notably, the model inclusion of these three widely separated timescale *requires* approximately 24 hours of simulation time to reproduce the critical emergent dynamics of AF progression, and to our knowledge, these simulations represents the *longest duration simulations* reported of beat-to-beat cardiac electrophysiology tissue dynamics.

A simplified tissue representation represents a final important assumption of the model. To isolate the role of calcium feedback regulation and the remodeling processes, we assume an isotropic, 2D geometry with insulating boundaries, with a surface area approximating the total area of both atria [30]. Voltage and calcium activation are known to be spatially variable in the atria [46]. Previous studies have simulated heterogeneous tissue using specific static distributions of ion channel conductances or intercellular coupling, representing either healthy or pathological states [47] Additionally, populations of models approaches, typically performed in single cell studies, have investigated responses for distributions of ion channel conductance levels [48–50]. However, in this study, since the ion channel conductances and tissue properties are dynamic and are ultimately governed by the feedback mechanism, initial variation in the conductance values will converge to the level determined by the particular *Ca*_*tgt*_ and the cell’s activation rate, as shown in Figure S6. Therefore, for simplicity, we define the initial conditions for the conductance values with a homogeneous value of 1 for all *γ*, i.e., the baseline model values, and we assume that tissue heterogeneity arises from heterogeneous *Ca*_*tgt*_ values, as shown in Figure 3E.

In addition, the atrial anatomy is more complex: the left atrium is spheroidal, while the right atrium is cylindrical. They are joined together by the interatrial septum, and the auricles of both atria have a trabeculated, network like structure. The atrial wall further has multiple layers [51], and conduction in tissue is anisotropic [52]. Further, these structures remodel during the progression of AF, leading to perturbed conduction: atrial surface area increases, and the tissue becomes fibrotic [30] The development of fibrosis during the course of AF is triggered by fast pacing, but also depends on local inflammation [53] and mechanical stretch [54]. For simplicity, we represent these structural changes as the dynamic intercellular coupling factor *γ*_*D*_, which summarizes the long term changes in tissue conduction. All of these details complicate the atrial geometry (fibrosis in particular providing distinct anchor points for re-entrant waves) leading to an increased possibility of re-entry.

Therefore, in order to separate the effects of complex structure and remodeling, we demonstrate that the simplified geometry, coupled with heterogeneous cells and intercellular connections, is sufficient to capture the initiation and progression of AF-associated re-entrant episodes.

### Ion channel regulation and adaptation to pacing

We highlight that the single cell model integrating the two assumptions discussed above (a single *Ca*_*tgt*_ and expression level correlations) qualitatively reproduces the key response of atrial cells to rapid pacing (Figure 1A-D): ion channel expression changes in response to pacing, leading to the characteristic reduction in the CaT amplitude, APD, and resting membrane potential, as well ultimately to a homeostatic prevention of *Ca*_*i*_ overload [43, 55]. The simulated rapidly paced atrial cell further establishes a natural, emergent steady state of remodeled expression levels, similar to experimental data, and reproduces the reversibility of pacing-induced changes [42], as seen in Figure 1A. In this context, it is also interesting to note that chronic bradycardia (in the ventricle) also leads to electrical remodeling, albeit in the *opposite* direction, leading to a decrease in *I*_*Kr*_ and *I*_*Ks*_, and a faster activation of *I*_*CaL*_, driving a corresponding increase in APD [56]. This phenomenon could also likely be explained by this model, demonstrating the broad utility of this simplified calcium regulatory feedback model framework: a slower pacing rate would naturally lead to a shift in expression to the *left* of the graph in Figure 1E, which is also equivalent to the 1000 ms baseline pacing in a cell with higher *Ca*_*tgt*_, as shown in Figure S10.

### Emergent spiral wave stabilization

By extending the single cell model to 2D tissue, we investigate for the first time the behavior of re-entrant waves in the presence of dynamic remodeling of the underlying substrate. In homogeneous tissue (defined as uniform *Ca*_*tgt*_), fast pacing drives changes in the tissue, leading to a transition from unstable spiral trajectories to the formation of a quasi-stable core.

It has previously been shown that defining parameters in an atrial cell model to correspond to changes seen in AF remodeling leads to the stabilization of spiral meandering (e.g. [16, 57]). Notably, these studies consider the underlying tissue to be static. Recently, Jaeger et al [58] used a dynamic model of ion channel expression, in which the conductance of *I*_*CaL*_ is variable. Their single cell results are in accordance with the results we report in Figure 1. Further, they show that *I*_*CaL*_ conductance values predicted by fast pacing in the single cell can induce a sustained re-entry wave if used in a ring geometry (equivalent to a pulmonary vein). However, their tissue simulations use static values of the *I*_*CaL*_ conductance, taken as snapshots from different time points in the single cell simulations. Therefore, the tissue substrate is not directly coupled to the electrical activity it sustains and, as in previous work, is comparable to a study with constant parameters. Further, more broadly, prior work in other systems with significantly different timescales (such as adaptive networks) that treat the slow modifying variables as constants lead to qualitatively different results when compared to simulating the complete coupled system [59].

In our 2D simulations, the electrical activity is coupled at all times (through *Ca*_*i*_) to the dynamic remodeling of the tissue substrate, which enables a series of novel observations. As noted above, fast pacing leads to the stabilization of spiral wave trajectories. The properties and specific patterns of re-entrant wave trajectories are known to be affected by the balance between tissue excitability and the excitation threshold [60], as well as by the tissue APD and conduction velocity restitution curves [61]. All of these properties are dynamically regulated in our model. Thus, rapid pacing leads to a dynamic transition to a new stable regime with lower excitability, shorter APD, and slower conduction velocity, which collectively in effect, constitute a trade-off between maintaining calcium homeostasis at the single cell level for increased susceptibility to stable spirals at the tissue level.

Notably, in the setting of the uniform substrate, the location of the stable spiral core itself leads to an interesting spatial pattern, as noted above: due to lower activation close to the spiral tip trajectory, the average *Ca*_*i*_ of the cells in this spatial region is lower, leading to an area of comparatively less remodeled tissue (i.e. *γ* values closer to the baseline of 1, see Figure 2B), forming a ‘ring’ shape. This pattern introduces an inhomogeneity in the tissue, which would otherwise be completely uniform. The established spiral wave then paces the entire tissue at a rate that is fast enough to maintain a new steady state in the electrical remodeling (see Figure S8, for *t >* 1 hour), and crucially one that is favorable for spiral wave stability. However, over the longest timeline of approximately 24 hours, we also observe that the new state is not static: there is a long term drift of the spiral core, as well as a transition in the re-entrant pattern. Finally, it is also interesting to note that remodeling can act as an anchor point for induced activity. This is apparent in the low *Ca*_*tgt*_ tissue highlighted in Movie S2. A late spiral, induced in the middle of the tissue, ‘jumps’ and anchors to a ring remodeled by a previous re-entrant wave (at approximately the 22 hours mark). The implications of these long-term trends are discussed in more detail below.

### AF initiation and progression

We lastly consider the response to rapid pacing in a large, heterogeneous atrial tissue (Figures 3 - 6). First, we demonstrate that rapid pacing alone induces re-entry, as shown in the earliest studies of AF progression [2]. As observed in Movie S5, different regions within the heterogeneous tissue respond differently to fast pacing: calcium transient abnormalities lead to a localized, temporary, conduction block around which a first re-entrant wave forms. This pattern is in accordance with experimental data showing irregular calcium signaling preceding re-entry [62].

The initially induced re-entry is however unstable, and subsequent pacing induces further transient re-entrant episodes. These in turn continue to contribute to substrate remodeling, such that re-entrant episodes become progressively longer (Figure 3B). This is one of the key results of our study, that the well-known dynamics of AF progression, seen in both experimental models (e.g., the cornerstone example study of Wijffels et al [2]) and patients [5], are an emergent behavior of the fully integrated atrial tissue model. As remodeling progresses, the tissue can sustain more re-entrant waves (Figure 5B), leading to more complex electric activity, consistent with studies showing an increase in the number of rotor locations in patients with longer history of AF [63]. The same phenomenon is also illustrated by Climent et al [20], which studied the evolution of re-entry in cell cultures: the increased number in spiral tips (or phase singularities) and the resulting fragmentation of the re-entrant waves is directly comparable with the our findings in Figure 5E, F. These changes in turn modify tissue voltage activity: simulations also capture the increase in the dominant frequency of activation (Figure 5B, cf. Martins et al [16]), as well as the fragmentation of the pseudo-ECG, indicative of more complex activation patterns (Figure 5C, cf. Wijffels et al [2], Figure 4). It is also interesting to note that the dynamics of both the spiral wave count and the dominant frequency are biphasic. There is an initial rapid increase, which corresponds with the timescale of electrical remodeling, followed by a more gradual increase as intercellular remodeling progresses (see *γ*_*CaL*_ and *γ*_*D*_ in Figure 3C, D, respectively). The initial electrical changes are sufficient to induce re-entry and thus transient AF episodes (corresponding to paroxysmal AF), while the longer term intercellular changes are necessary for the transition to permanent re-entry. This timeline is consistent with the concept of a “second factor” in AF progression, in which longer term structural changes are necessary for the transition to permanent AF [64, 65]. Finally, we also observe instances (i.e., particular *Ca*_*tgt*_ spatial patterns) in which the atrial tissue dynamics does not reach a persistent phase (Figure S16), and rather exhibit behavior resembling long term paroxysmal AF.

The characteristics of electrical and calcium activity in the remodeled tissue (i.e., at the end of the simulation from Figures 3 - 6) also match published experimental data associated with AF remodeling. The spatial pattern of APD is more heterogeneous than in the corresponding tissue paced at a baseline of 1000 ms (Figure S13, C), corresponding to published data [66]. Notably, the spatial locations corresponding with the transition between high and low APDs correspond with where re-entry is initiated (Movie S5) and where the spiral tips tend to localize (Figure 5 D), equivalent to data published by Avula et al [67]. Further, conduction velocity is slower and more irregular (Figure S13, D, cf. [68, 69]). Voltage and calcium activity are dissociated (Figure 6 E), hallmarks of highly remodeled atrial tissue [70], and the calcium activity exhibit an interesting, complex pattern of alternating high and low peaks, reminiscent of recent results from Iravanian et al [71] that show complex periodicities in ventricular fibrillation. Notably, the only constraints we impose are the heterogeneous *Ca*_*tgt*_ distribution and aforementioned cellular level assumptions. Therefore, all of the results in heterogeneous tissue and their agreement with experimental data are emergent, and moreover provide a mechanistic explanation for the phenomena discussed.

The long-term dynamics of the re-entrant waves observed present novel consideration. In the homogeneous tissue with the baseline *Ca*_*tgt*_ = 258 *nM*, a single spiral wave stabilizes. On a short timescale, the location can appear to be stationary. However, as discussed above, we observe a long term drift (Figure S8), along with a dynamic spiral tip pattern (Movie S1). In effect, we identify a novel spiral drift mechanism (for a review of known causes for drift see [72]) caused by substrate remodeling. This mechanism is directly influenced by the *Ca*_*tgt*_ value: a high value (*Ca*_*tgt*_ = 320 *nM*) prevents spiral wave formation altogether (Movie S3), while a low value (*Ca*_*tgt*_ = 200 *nM*) leads to dynamic activity, including shifting spiral locations, along with a complex pattern of remodeling (Movie S2). The heterogeneous tissue *Ca*_*tgt*_ values span the interval between these two extreme levels; it is thus notable that we find the highest density of rotor tips in the transition areas between high and low *Ca*_*tgt*_, where the homogeneous simulations would predict the most stable re-entry.

Early re-entrant activity can exist without a single preferential rotor location, resembling the earliest AF mathematical modeling studies [73, 74], where Moe et al observed multiple chaotic wavelets. However, in the example shown in Figures 3 - 6 preferential rotor locations appear in the low *Ca*_*tgt*_ areas, especially towards the end of the simulation, pointing to a late organization of activity. We observe this pattern of spatially consistent re-entrant hotspots in other examples of permanent re-entry (see Movies S8 and S11), in which a noticeable spatial remodeling in the same ‘bump’ pattern is induced, as in the homogeneous case. This leads to the intriguing hypothesis that AF driver regions and their associated remodeling pattern [75] are inherently dynamic and can self organize, induced by spiral waves forming their own substrate niche. Overall, our model predicts situations in which both chaotic wavelets and more stable rotors [76] can exist. An interesting next step is to delineate conditions necessary for either case, as well as the corresponding clinical or experimental situations.

### Limitations and future directions

Our model is limited by the large computational requirements necessary to simulate the long periods of time imposed by the timescales simulated. Therefore, we adopted a relatively simple geometry, as discussed above. Atrial fibrillation has been studied in computational models with increasingly more realistic geometries, incorporating whole atria 3D geometries derived directly from individual patient data [22]. These models require significant spatial resolution leading to high computational costs and, as a consequence, simulation studies generally only reproduce at most several second of activity. Therefore, simulations only replicate snapshots of healthy or pathological tissue, instead of a dynamic continuum. However, it would be an interesting next step to integrate more complex aspects of atrial anatomy with our model, such as the network topology of the trabeculate auricles, periodic boundaries to mimic the cylindrical or spherical shapes of the atria, as well as multiple tissue layers to replicate the endo- and epicardium [51].

The specific electrophysiological model used, the Courtemanche et al model [24], introduces additional inherent limitations in the study. First, this model does not include important atrial-specific ion channels such as the two-pore-domain or the small-conductance Ca^2+^-activated potassium channels, which have been shown to change in AF and are described in more complex recent models [77]. Further, the Courtemanche model does not exhibit early or delayed afterdepolarizations (EADs or DADs) [57], which have also been proposed as a source for AF initiation and maintenance alongside re-entrant waves, especially as the origin of rapid impulses from the pulmonary veins [78]. DADs are known to be caused by spontaneous calcium releases, in particular during calcium overload, which in turn depend on localized calcium release units (i.e., t-tubules and sarcoplasmic reticulum). Recent models have succeeded in providing detailed descriptions of these subcellular structures and their pathological disruption [79, 80]. However, these advanced models greatly increase computational complexity - the simulation of a single cell becomes comparable (in resource requirements) to a 2D tissue using a simpler model.

However, despite these limitations, the Courtemanche model continues to be used in large scale tissue simulations, for which computational efficiency is an important requirement. For example, it serves as a basis for a recent study by Boyle et al[81], in which the authors use whole-atria simulations to incorporate patient-specific data and successfully predict personalized arrhythmia risk and re-entrant wave locations. Thus, while more detailed and complex models have since been developed, the Courtemanche model still provides a sufficiently accurate approximation of atrial electrophysiology and arrhythmic activity. In our case, as with whole-chamber or subcellular simulations, the computational cost is significant due to the multiple timescales and simulation durations considered. Therefore, we chose the Courtemanche model to represent atrial ionic current dynamics as a compromise between computational cost and complexity. Despite its relatively simple (but therefore efficient) formulation, it still describes detailed calcium cycling dynamics and most of the currents known to remodel in AF, and therefore the APD, CV, and excitability changes needed to model re-entrant wave activity. Finally, we note that the feedback model consists of a simple set of equations, which could easily be adapted to the more complex models mentioned above, and indeed would be a natural extension of the current work.

More generally, as AF arises naturally in patients without outside interventions such as pacing, it is highly likely that there are other long-term substrate changes, which could be modeled by modifying the inherent properties of the feedback mechanism, such as the *Ca*_*tgt*_ or the channel co-expression ratios. Such underlying changes could, for example, be the cause of rapid impulse generation at the pulmonary veins, which would lead to a naturally occurring regime of intermittent fast pacing driving further remodeling. A similar approach could also be readily applied more broadly to other settings of pathological long-term remodeling, such as in the ventricles during heart failure. In conclusion, we have introduced a novel framework to uncover the mechanisms underlying the full progression of AF, developing novel methodologies to enable the first ever simulations of this long timescale and spatiotemporally complex pathological process.

### Competency In Medical Knowledge

Atrial fibrillation is the most common cardiac arrhythmia. It is notable for being a progressive disease, with pathological processes that span a wide timescale: millisecond changes in ion channel and voltage dynamics driving protein expression and structural changes over days or weeks. Our study proposes a unifying mechanism driving AF progression, based on long-term calcium homeostasis. Simulations predict spontaneous spiral wave stabilization and progressively longer re-entrant electrical episodes that eventually become permanent, hallmarks of the disease. Critically, we observe a new phenomenon: reentrant waves, through long-term remodeling, pattern their own emergent tissue ‘niches’ that ensure long-term stability of arrhythmic activity.

### Translational Outlook

Our study unifies different lines of research (ion channel co-regulation, calcium homeostasis, AF natural history) into a single framework, and provides for the first time a computational investigation of the entire progression (i.e., initiation to remodeling to stabilization) of AF. This opens a new perspective on in-silico testing of novel drugs or drug combinations that can directly address AF progression, leading to the identification of new disease-modifying treatment strategies in AF.

