## Supplemental Methods, Tables, and Figures for "Calcium homeostatic feedback control predicts atrial fibrillation initiation, remodeling, and progression"

### 1 Supplementary Methods

#### 1.1 Model

**Atrial myocyte electrophysiology and calcium regulatory feedback model**

The full atrial model integrating excitation-transcription regulation with the electrical-calcium dynamics is based on the Courtemanche human atrial ionic model [1], coupled with the calcium regulatory feedback model originally proposed for neurons by O'Leary et al [2]. The mathematical formulation of the model is given by:

$$\tau_x \frac{dm_x}{dt} = Ca_{tgt} - Ca_i, \tag{S1}$$

$$\tau_\gamma \frac{d\gamma_x}{dt} = m_x - \gamma_x, \tag{S2}$$

where $Ca_{tgt}$ is the homeostatic calcium target in a given atrial myocyte, $Ca_i$ is the intracellular calcium concentration, $m_x$ represents a normalized mRNA expression (relative to baseline), and $\gamma_x$ is a scaling factor for ionic current conductances.

The regulatory feedback model is coupled to the atrial cell electrophysiology model through scaling factors $\gamma_x$, where $x \in X = \{Na, CaL, ito, Kur, K1, Ks, RyR_{leak}, SERCA, NCX\}$, such that the conductance for current $I_x$ or flux $J_x$ is given by $\gamma_x g_x$, where $g_x$ is the baseline conductance given in Courtemanche et al [1]. The specific currents

and fluxes regulated by the calcium feedback model are based on available data in Grandi et al. [3], as well as those with representation in the Courtemanche model.

In a single atrial myocyte, the transmembrane voltage ($V$) dynamics are represented succinctly as

$$\frac{dV}{dt} = -I_{ion}(V, Ca_i, \boldsymbol{g}, \boldsymbol{\gamma_x^V})/C_m, \tag{S3}$$

where $C_m$ is the membrane capacitance and the sum of ionic currents $I_{ion}$ depends on $V$, intracellular calcium $Ca_i$ (via $Ca_i$-dependent ionic currents), vector $\boldsymbol{g}$ consisting of all ionic model gating variables [1], and vector $\boldsymbol{\gamma_x^V}$ consisting of scaling factors for all voltage-dependent currents coupled to the feedback model: $I_{Na}, I_{CaL}, I_{NCX}, I_{ito}, I_{Kur}, I_{K1}$, and $I_{Ks}$.

Similarly, the intracellular calcium ($Ca_i$) dynamics can be written succinctly as

$$\frac{dCa_i}{dt} = f(V, Ca_i, \boldsymbol{\gamma_x^{Ca_i}}), \tag{S4}$$

where the function $f$ depends on $V$ (via voltage-dependent calcium currents), and vector $\boldsymbol{\gamma_x^{Ca_i}}$ consisting of scaling factors for the intracellular calcium fluxes and transmembrane calcium currents coupled to the feedback model, $J_{RyR_{leak}}, J_{SERCA}$ and $I_{CaL}, I_{NCX}$, respectively. A complete description of the electrophysiological model can be found in Courtemanche et al [1].

#### Estimation of τ values governing current / flux remodeling dynamics

Here, we illustrate the methodology to estimate the time constants for AF remodeling, based on experimental values. We define the scaling factors for the healthy state as $\gamma_x^0$, from which it follows that $\gamma_x^0 = 1$ for $\forall x \in X$, and define the value in chronic AF as $\gamma_x^{AF}$.

As described above, $\gamma_x$ obeys the calcium feedback differential equations:

$$\tau_x \frac{dm_x}{dt} = Ca_{tgt} - Ca_i, \tag{S5}$$

$$\tau_\gamma \frac{d\gamma_x}{dt} = m_x - \gamma_x, \tag{S6}$$

where $m_x$ is a normalized mRNA scaling factor, $\tau_x$ is a time constant for each $x \in X$ (with units of concentration · time), and $\tau_\gamma$ is a time constant representing the timescale of mRNA expression, common to all $x$ (with units of time). Note that $\tau_x$ governs the response of each scaling factor to the $Ca_i$ feedback.

We note that, from Eq. S5, for any two species $(x, y)$

$$\tau_x \frac{dm_x}{dt} = \tau_y \frac{dm_y}{dt}. \tag{S7}$$

Integrating both sides with respect to time results in the following:

$$\tau_x \int_{t_0}^{t_1} \frac{dm_x}{dt} dt' = \tau_y \int_{t_0}^{t_1} \frac{dm_y}{dt} dt', \tag{S8}$$

such that

$$\tau_x (m_x(t_1) - m_x(t_0)) = \tau_y (m_y(t_1) - m_y(t_0)). \tag{S9}$$

Additionally, at steady state, we note that,

$$m_x = \gamma_x, \tag{S10}$$

i.e., the normalized mRNA expression and ionic conductance scaling factor are equivalent. We substitute $m_x$ from Eq. S10 in Eq. S9 and obtain

$$\tau_x (\gamma_x(t_1) - \gamma_x(t_0)) = \tau_y (\gamma_y(t_1) - \gamma_y(t_0)). \tag{S11}$$

Further, we take $\gamma_x(t_1) = \gamma_x^{AF}$ (the value in chronic AF), $\gamma_x(t_0) = \gamma_x^0$ (the value in the healthy state), and define species $y$ similarly, such that

$$\frac{\tau_x}{\tau_y} = \frac{\gamma_y^{AF} - \gamma_y^0}{\gamma_x^{AF} - \gamma_x^0}. \tag{S12}$$

Note that this relationship holds, assuming the healthy state and the chronic AF state are at equilibrium, according to Eq. S10. This assumption is verified by the single cell simulations in Figures 1, S2-S3. Finally, we set $\tau_y = \tau_{Na} = 150 \cdot 10^4$ nM $\cdot$ ms (in order to approximate the remodeling dynamics timescale from Goette et al[4]), which together with the $\gamma_x^{AF}$ values derived from Grandi et al. [3] (given in Table S1) enable us to obtain all other $\tau_x$ values relative to $\tau_{Na}$. Note that $\tau_{Na}$ and $\tau_\gamma = 2 \cdot 10^4$ ms collectively set the overall timescale of the feedback system. The values we obtain for all $x$ are given in Table S2.

To simplify the numerical calculations, based on recent demonstration by Jaeger et al [5] that there is a sufficient separation of timescales, such that Eqs. S5-S6 can be consider to be in quasi-steady state relative to the faster timescale of the ionic currents and electrical dynamics, then Eq. S11 holds for any $t$, such that

$$\gamma_x(t) = \frac{\tau_{Na}}{\tau_x}\left(\gamma_{Na}(t) - \gamma_{Na}^0\right) + \gamma_x^0, \tag{S13}$$

which, since $\tau_x$ and $\tau_{Na}$ are constants, is a simple linear relationship. Using the $\tau_x$ values as derived above allows us to directly calculate the corresponding $\gamma_x(t)$ value from the time-dependent $\gamma_{Na}(t)$ (for all $x$ except $Na$), thus eliminating the need to compute 8 out of the 9 ordinary differential equation pairs (Eqs. 1-2) governing the feedback model.

#### Atrial tissue model

We extend the single atrial myocyte model to simulate a two-dimensional tissue by the partial differential reaction-diffusion equation governing $V$,

$$\frac{\partial V}{\partial t} = \nabla \cdot [(\gamma_D D)\nabla V] - I_{ion}(V, Ca_i, \boldsymbol{g}, \boldsymbol{\gamma_x^V})/C_m, \tag{S14}$$

where $D$ is the baseline diffusion coefficient and $\gamma_D$ is an isotropic local scaling factor representing remodeling of intercellular coupling. Note that intracellular calcium and the calcium feedback model are thus similarly governing 'local' dynamics, with regulation occurring at each spatial location within the atrial tissue, indirectly coupled via the 'global' tissue-scale electrical dynamics.

Intracellular coupling scaling factor $\gamma_D$ is similarly governed by the calcium regulatory feedback model, as with the ionic current and flux conductances:

$$\tau_D \frac{dm_D}{dt} = Ca_{tgt} - Ca_i, \tag{S15}$$

$$\tau_{\gamma_D} \frac{d\gamma_D}{dt} = m_D - \gamma_D, \tag{S16}$$

Importantly, to account for the differences in timescales between ionic current and intracellular coupling remodeling, we assume that intercellular coupling changes are several orders of magnitude slower than electrical remodeling, and thus take $\tau_{\gamma_D} = 1000 \cdot \tau_\gamma$, with the specific values given in Table S2.

#### Timescales of the feedback model

There are three timescales in the integrated electrophysiology-calcium regulatory feedback model: beat-to-beat action potential and calcium transient dynamics governed by the Courtemanche human atrial model [1], electrical remodeling, governed by parameters $\tau_\gamma$ and $\tau_{Na}$, and intercellular coupling remodeling, governed by $\tau_{\gamma_D}$ and $\tau_D$.

Two foundational studies demonstrated that the electrical remodeling occurs on a timescale of 1 to 24 hours [4, 6]. Both studies quantified this timescale by analyzing changes in the effective refractory period. Goette et al [4] found that most of the effect of electrical remodeling occurred during the first hour of rapid pacing in the dog atria. Wijffels et al [6] found that the effective refractory period reached a minimum after 24 hours of rapid pacing in the goat atria. For the sake of computational efficiency, in our study, parameters $\tau_\gamma$ and $\tau_{Na}$ were chosen to reflect the faster timescale, such that the electrical remodeling reaches steady state after around one hour (c.f., Figures 1, 2). However, due to the large separation of timescales between action potential dynamics (milliseconds) and electrical remodeling (hours), it is reasonable to consider that the results observed in this study to be relevant for the longer electrical remodeling timescales as well.

Intercellular coupling (or structural) remodeling is known to occur on an even slower timescale than electrical remodeling [7, Table 1], on the order of weeks or even months after AF initiation. To represent intercellular coupling remodeling in the model, while accounting for the large computational cost and maintaining a reasonable simulation time, we assume that this remodeling is significantly slower than electrical remodeling, and therefore choose $\tau_{\gamma_D} = 1000 \cdot \tau_\gamma$. Finally, we choose $\tau_D$ so that the scaling factor value $\gamma_D$ during rapid pacing and re-entrant activity varies from 0.4 to 0.8 (e.g., Figure 3), relative to the baseline of 1, consistent with the degree of reduced coupling observed [8, 9].

#### Spiral tip tracking, pseudo-ECG calculation and APD calculation

Following the method described by Aron et al [10], we detect the presence of spiral waves by identifying phase singularities throughout the tissue. We compute the phase $\phi$ associated at each spatial point $(x, y)$ in the 2D grid:

$$\phi = atan2(V_{norm} - V_0, h - h_0),$$

where $atan2$ is the 2-argument arctangent function, $V_{norm} = (V - V_r)/V_{amp}$ is the normalized transmembrane voltage, $V_r = -90$ mV, $V_{amp} = 100$ mV, $h$ is the $I_{Na}$ inactivation gating variable, and $V_0 = 0.3$, $h_0 = 0.4$ are reference points along the action potential phase. Next, we determine the respective trigonometrical quadrant for all 8 neighbors of each particular spatial location. If at least one neighbor is in each of the four quadrants, we identify the point $(x, y)$ as a phase singularity.

Additionally, to represent the summation of the atrial tissue electrical activity, a pseudo-ECG, $\bar{V}$, is calculated by the spatial average of $V_{norm}$, across all spatial locations.

In order to compute APD, we normalized voltage values such that the resting membrane potential is 0 and the peak of activation is 1. APD was defined as the duration between the start of the activation and the return to a normalized value of 0.2. In single cell simulations (Figure 1, S2, S3, S4) the cells were paced at the respective cycle length for 5 s, and the APD was computed for the last activation. In the 2D tissue measurement (Figure S13), the tissue was paced at a cycle length of 1000 ms for 10 s and APD was calculated for the last beat, for all ($1024 \times 1024$) spatial locations in tissue.

#### Tissue pacing protocols

Rapid pacing in the atrial tissue is performed at a frequency of 10 Hz, by resetting $V$ to 0 mV every 100 ms at all tissue spatial locations within 0.625 cm from $x = 0$ and $y = 0$ (i.e., all locations within the quarter circle with radius 0.625 cm centered on the origin, which is located in the lower left corner of the tissue). In the homogeneous tissue simulations (Figures 2-S8, S9-S11), rapid pacing is performed for an interval of 10 s, after which a spiral wave is initiated by cross-stimulation. Every 10 ms, the absence of electrical activity throughout the tissue is assessed, as determined by $\bar{V} < 0.1$. If electrical activity is extinguished, then the tissue is paced for another 10 s interval followed by cross-stimulation.

In heterogeneous tissue (Figures 3-6, S15), rapid pacing in the atrial tissue is similarly performed at a frequency of 10 Hz, by resetting $V$ to 0 mV every 100 ms at all tissue spatial locations within 0.625 cm from $x = 0$ and $y = 0$; however cross-stimulation is not applied. Every 1 ms, the presence of phase singularities, representing the presence of at least one spiral wave (as described above) is assessed. If such a phase singularity is detected, the pacing is continued for a further 1.5 s (15 cycles), after which pacing is stopped. The additional 15 cycles of pacing after the appearance of phase singularities are performed in order to obtain consistent re-entrant activity.

#### Heterogeneous $Ca_{tgt}$ maps

The heterogeneous $Ca_{tgt}$ maps used in Figures 3-6, S15 were generated by smoothing uniform random noise using a diffusion kernel, using an approach similar to our previous work generating random maps [11]. After smoothing, the random maps were normalized, such that $Ca_{tgt}$ values in the random map vary over the range from [200, 320] nM. Using this approach, the strength of the random map smoothing determines the relative size of the 'features,' where the features are the different regions of high and low $Ca_{tgt}$ values. All $Ca_{tgt}$ maps used in the simulations are shown in Figure S15.

### 1.2 Numerical methods

All numerical simulations were integrated using forward Euler's method with time step $\Delta t = 0.1$ ms. For 2D tissue simulations, we simulate homogeneous and heterogeneous tissues of size 6.4 cm x 6.4 cm (512 x 512) and 12.8 cm x 12.8 cm (1024 x 1024), respectively, using a spatial step $\Delta x = \Delta y = 0.0125$ cm, and discretizing the spatial derivative with an explicit nine-point stencil. The variable diffusion coefficient $D$ (due to time and spatial dynamics of $\gamma_D$) is locally approximated using the harmonic mean in order to ensure conservation of the voltage $V$, as shown in Shashkov et al.[12] and Patankar [13, p. 44-45]. All parameters used in the numerical simulations are given in Table S2.

In order to use the fastest possible time step ($\Delta t = 0.1$ ms), the Rush-Larsen method [14] was used to integrate the $m$ gating variable of the $I_{Na}$ current. Further, in the 2D simulations, we use an operator splitting scheme: the electrophysiolgical dynamics (i.e. $I_{ion}$ in Eq. S14) are calculated once for each overall timestep, while the diffusion operator is computed 10 times, with a $\Delta t_{diff} = \Delta t/10$, to take advantage of the computational efficiency of the explicit diffusion scheme while preserving numerical stability.

Single cell simulations were performed using MATLAB (Mathworks). The 2D tissue simulations were performed using CUDA on NVIDIA A100 GPUs through the Ohio Supercomputer Center [15] and the Delta system at the National Center for Supercomputing Applications through allocation MED230049 from the Advanced Cyberinfrastructure Coordination Ecosystem: Services and Support (ACCESS) program [16] (supported by National Science Foundation grants #2138259, #2138286, #2138307, #2137603, and #2138296).

All code used in this study is available at https://github.com/SHWeinberg/AFibProgression.

#### 1.3 Parameters

**Table S1.** Chronic AF remodeling scaling values derived from [3], Table 1.

| **Variable** | **Value** |
|---|---|
| $\gamma_{Na}^{AF}$ | 0.9 |
| $\gamma_{K1}^{AF}$ | 2 |
| $\gamma_{ito}^{AF}$ | 0.5 |
| $\gamma_{Kur}^{AF}$ | 0.5 |
| $\gamma_{Ks}^{AF}$ | 2 |
| $\gamma_{CaL}^{AF}$ | 0.5 |
| $\gamma_{RyR_{leak}}^{AF}$ | 2 |
| $\gamma_{SERCA}^{AF}$ | 0.75 |
| $\gamma_{NCX}^{AF}$ | 1.4 |

**Table S2.** Model parameters. ERP = effective refractory period.

| **Parameter** | **Value** | **Units** | Reference |
|---|---|---|---|
| $\tau_\gamma$ | $2 \cdot 10^4$ | ms | Estimated from ERP remodeling dynamics in [4] |
| $\tau_{\gamma D}$ | $2 \cdot 10^7$ | ms | Defined as significantly slower than electrical remodeling [7] |
| $\tau_{Na}$ | $9 \cdot 10^{-4}$ | nM · ms | Estimated from ERP remodeling dynamics in [4] |
| $\tau_{K1}$ | $-9 \cdot 10^{-5}$ | nM · ms | Derived from values in [3] |
| $\tau_{ito}$ | $1.8 \cdot 10^{-4}$ | nM · ms | Derived from values in [3] |
| $\tau_{Kur}$ | $1.8 \cdot 10^{-4}$ | nM · ms | Derived from values in [3] |
| $\tau_{Ks}$ | $-9 \cdot 10^{-5}$ | nM · ms | Derived from values in [3] |
| $\tau_{CaL}$ | $1.8 \cdot 10^{-4}$ | nM · ms | Derived from values in [3] |
| $\tau_{RyR_{leak}}$ | $-9 \cdot 10^{-5}$ | nM · ms | Derived from values in [3] |
| $\tau_{SERCA}$ | $3.6 \cdot 10^{-4}$ | nM · ms | Derived from values in [3] |
| $\tau_{NCX}$ | $-2.25 \cdot 10^{-4}$ | nM · ms | Derived from values in [3] |
| $\tau_D$ | $1.215 \cdot 10^{-4}$ | nM · ms | Derived from [8, 9] |
| $D$ | $1 \cdot 10^{-3}$ | $cm^2$/ms | [17] |
| $Ca_{tgt}$ | Single cell 258, Tissue [200,320] | nM | Calculated from time average $Ca_i$ in the atrial model [1] |
| $\Delta t$ | 0.1 | ms | - |
| $\Delta x$ | 0.0125 | cm | Length of a human atrial myocyte [18] |
| $x \times y$ | $6.4 \times 6.4$, $12.8 \times 12.8$ | $cm^2$ | Tissue size; derived from atrial surface values in [19], Table 2. |

### 2 Supplementary Figures

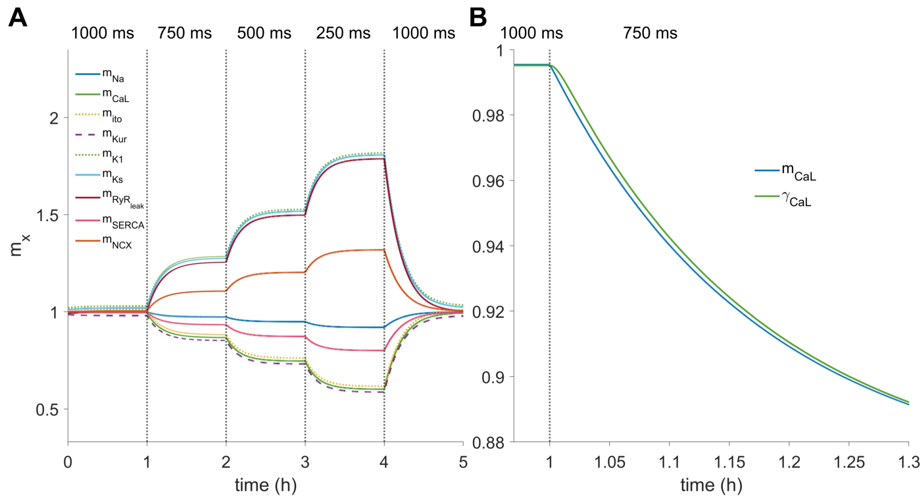

**Figure S1. A.** Normalized mRNA expression $m_x$ values for the simulation shown in Figure 1. **B.** Zoom-in of the transition from pacing at 1000 ms to 750 ms. Note that mRNA expression $m_{CaL}$ change occurs first, followed by conductance $\gamma_{CaL}$, which tracks closely in time.

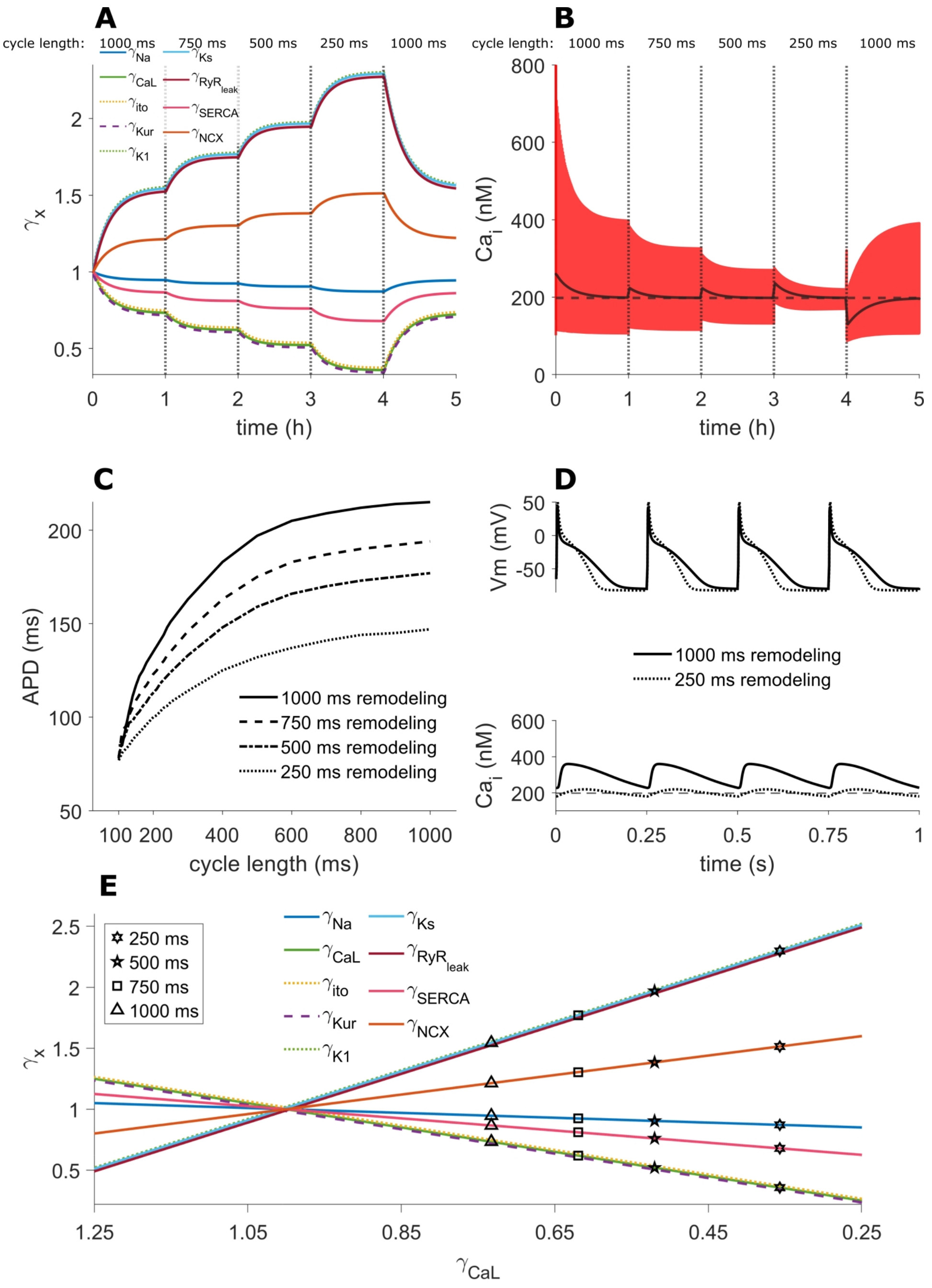

**Figure S2.** Electrical remodeling in the single cell with $Ca_{tgt} = 200$ nM. **A.** Remodeling at different cycle lengths. Note electrical remodeling is present at the baseline cycle length of 1000 ms, in addition to more pronounced remodeling as at faster pacing rates, compared to Figure 1 **B.** Calcium transients (CaTs, red) and moving average $Ca_i$ (black). **C.** APD restitution curves after different long term pacing rates. **D.** Electrical remodeling leads to shorter APDs and smaller CaTs. **E.** Linear relationship between conductances.

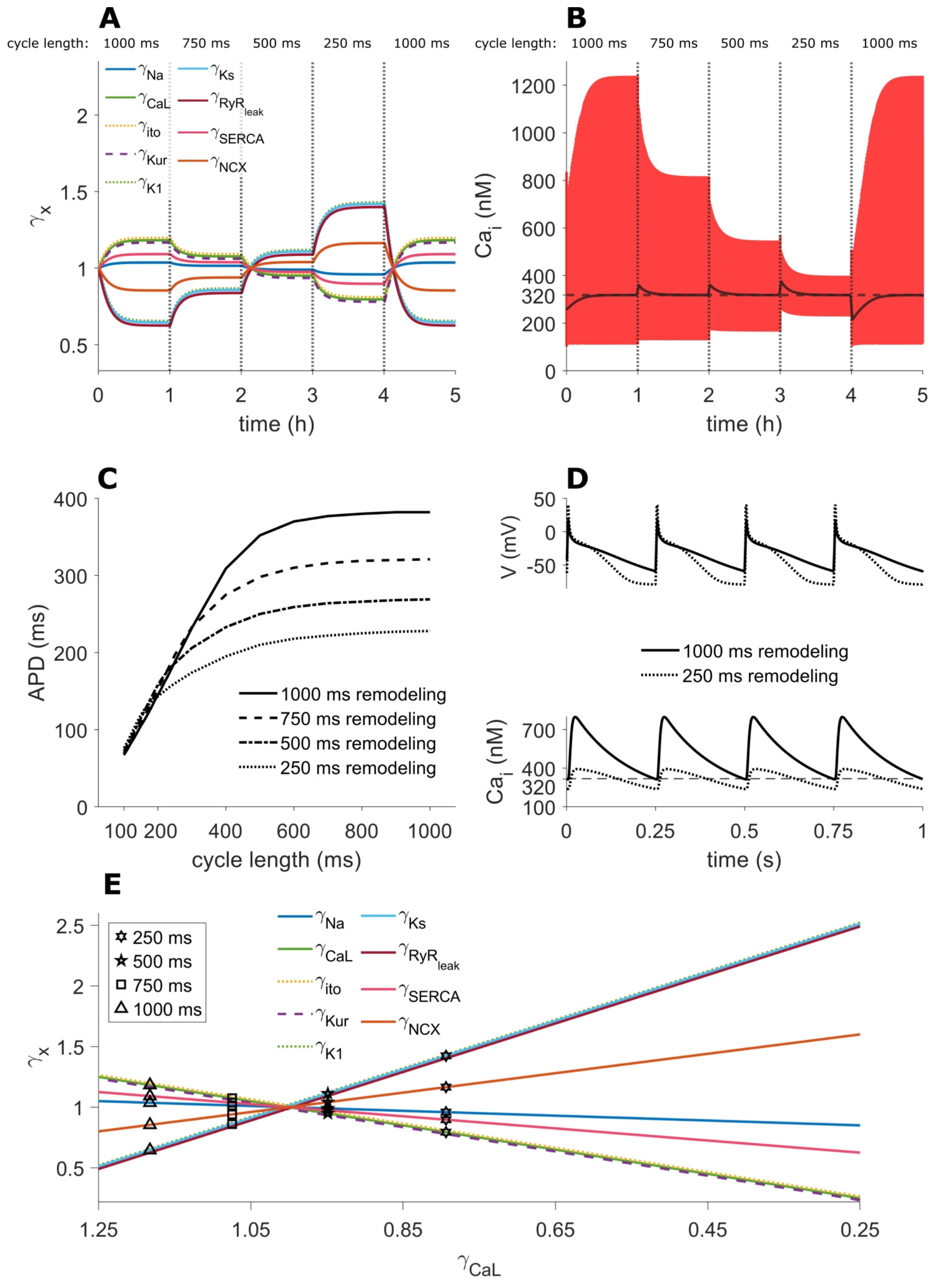

**Figure S3.** Electrical remodeling in the single cell with $Ca_{tgt} = 320$ nM. **A.** Remodeling at different cycle lengths. Note the reversed remodeling at the baseline cycle length of 1000 ms, and the decreased remodeling at fast pacing rates, compared to Figure 1. **B.** Calcium transients (CaTs, red) and moving average $Ca_i$ (black). **C.** APD restitution curves after different long term pacing rates. **D.** Electrical remodeling leads to shorter APDs and smaller CaTs. **E.** Linear relationship between conductances.

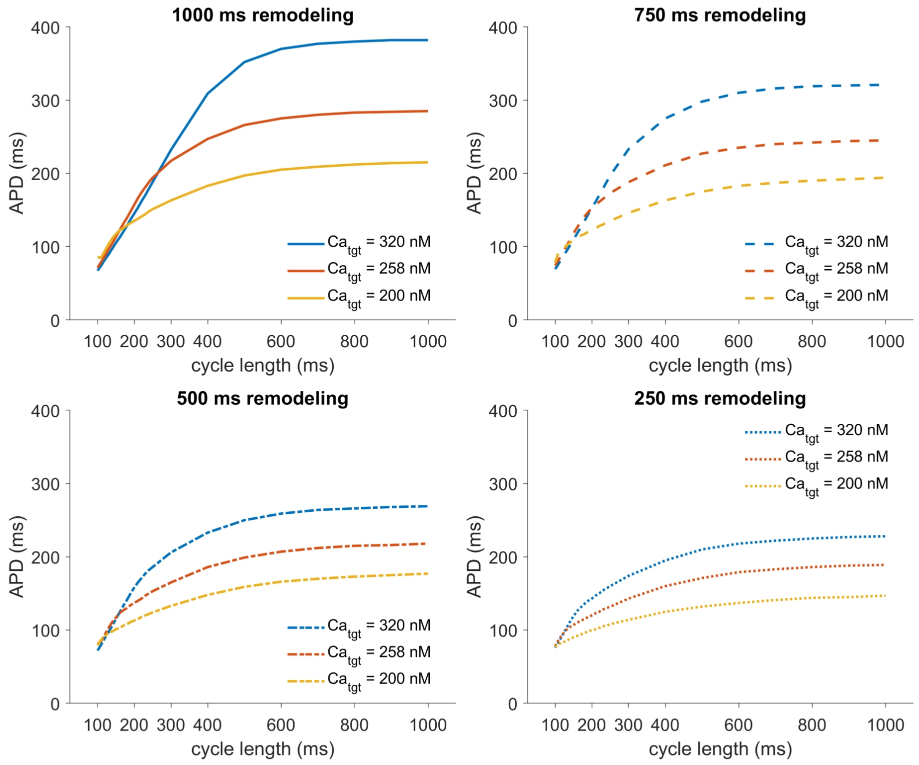

**Figure S4.** Action potential duration (APD) restitution curves after chronic pacing at different cycle lengths and $Ca_{tgt}$ levels. Note that APD restitution curves with higher $Ca_{tgt}$ are consistently closer to baseline at fast rates due to decreased electrical remodeling.

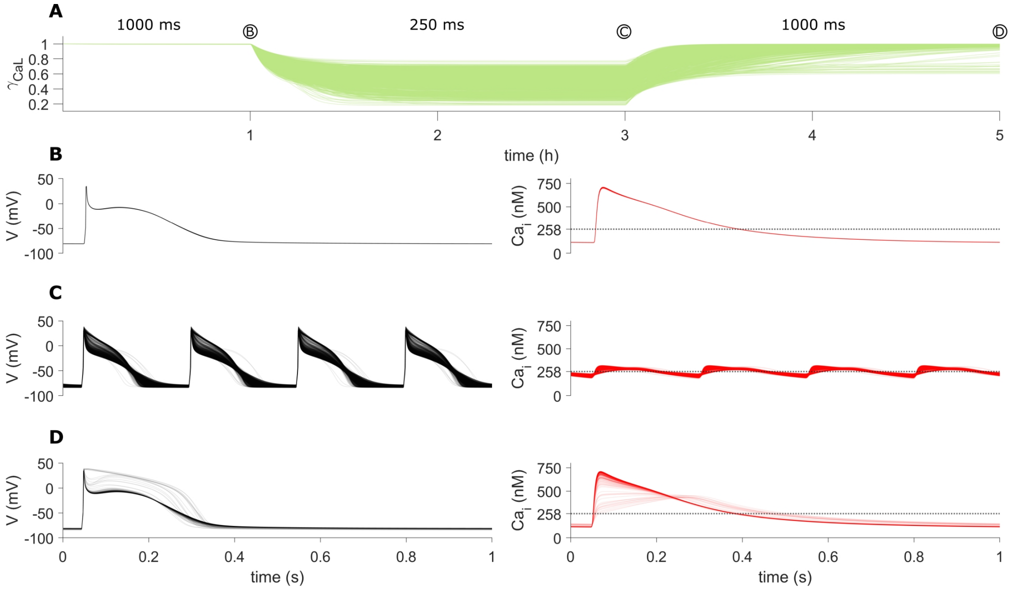

**Figure S5.** Single cell electrical remodeling with randomly varying remodeling time constants. Simulations are shown for 1000 random $\tau_x$ values, modifying the dynamics of each current conductance mediated by calcium feedback. Baseline values for each $\tau_x$ are scaled by a factor $2^k$, where $k$ is uniformly drawn from the interval $[-1, 1]$ Note that the linear relationship between conductances (highlighted in Figure 1) is still maintained, although with different slopes for each parameter set. **A.** Electrical remodeling $\gamma_{CaL}$ for all 1000 combinations of random $\tau_x$ values. Each cell approaches steady-state upon remodeling during rapid pacing at 250 ms, with average $Ca_i$ reaching $Ca_{tgt}$ (see B-D, right panels), highlighting robustness in the feedback model. Note that after the return to 1000 ms pacing, some conductances reach a new, intermediary steady state, suggesting a bi-stable behavior for some $\tau_x$ parameter sets. **B-D.** Voltage and $Ca_i$ traces at the highlighted time points. Note the variability in action potential morphology following rapid pacing (in panel C), some of which persists after the return to 1000 ms pacing (in panel D).

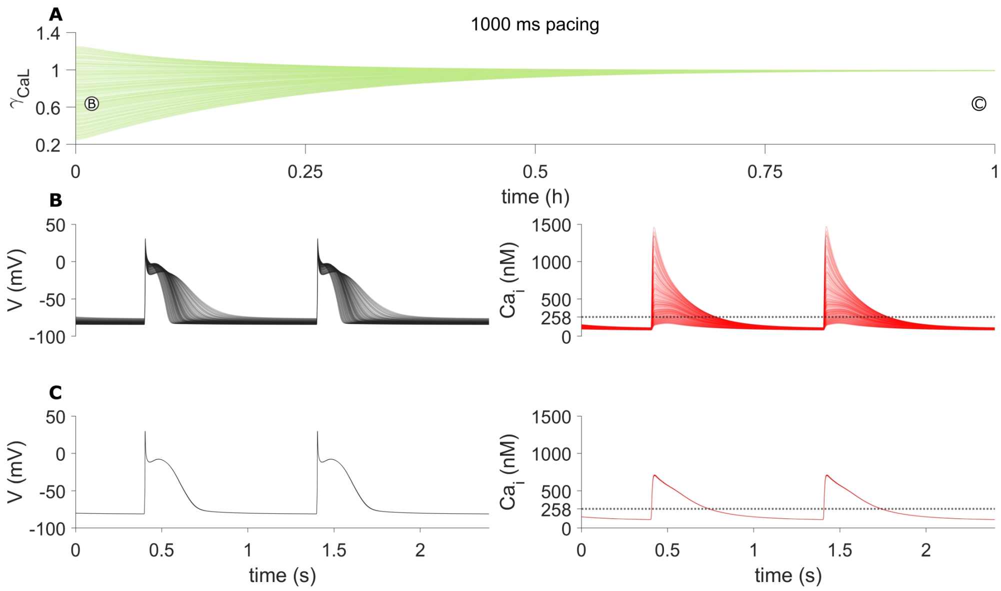

**Figure S6.** Single cell electrical simulation showing the evolution of 1000 different sets of initial conductance values, being paced at a 1000 ms cycle length. All cases have the same baseline $Ca_{tgt} = 258$ nM (shown as the horizontal dotted line in the left panels of B and C). All conductances are scaled between the extreme ranges from Figures S9 - S10. Note that the ratio between conductances is still maintained. **A.** The conductance values converge to the baseline fo 1 when paced at 1000 ms. **B.** Early activity shows a large degree of heterogeneity in both action potential and calcium transient shapes. **C.** Late activity is identical for all 1000 simulated cases, as conductances converge to the baseline of 1.

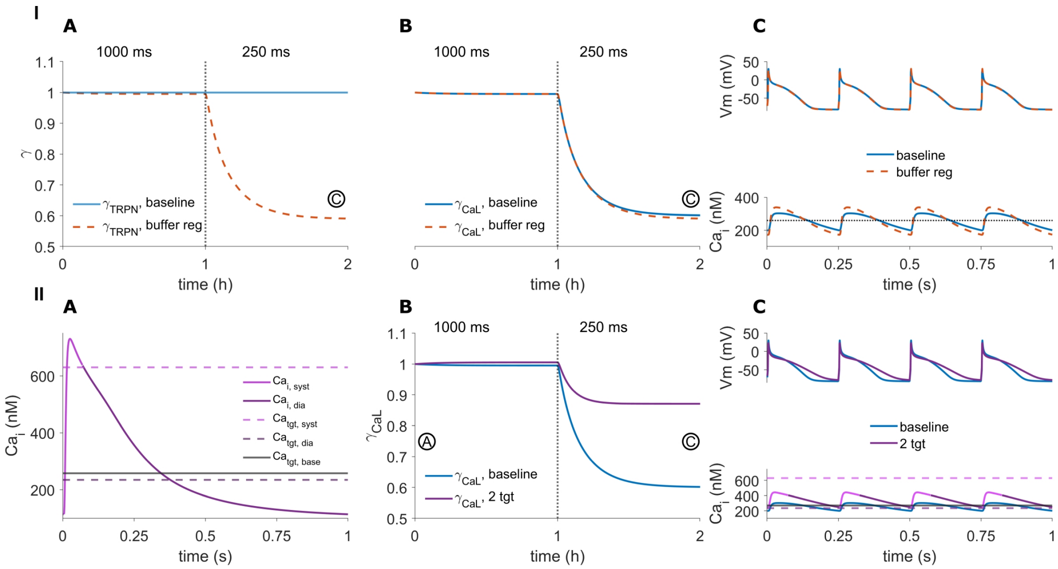

**Figure S7.** Different regulatory variants to the feedback model.

**I.** Simulations comparing the baseline model with a regulatory model that also governs the level of calcium buffers; the total troponin ($TRPN_{max}$), calmodulin ($CMDN_{max}$) and calsequestrin ($CSQN_{max}$) concentrations, as defined in the Courtemanche model, are also scaled by a corresponding $\gamma$ value controlled by the feedback model. **I-A.** $\gamma_{TRPN}$ decreases to half the healthy value at 250 ms pacing. $\gamma_{CMDN}$ and $\gamma_{CSQN}$ (not shown) have identical regulation. In the baseline model, these values are constant. **I-B.** Buffer regulation leads to minimal differences in ion channel expression compared to the baseline model, at both 1000 and 250 ms pacing. **I-C.** At 250 ms pacing, buffer regulation does not alter action potential shape, while the calcium transient is moderately shorter with a higher peak.

**II.** Simulations comparing the baseline model with a regulatory model that has two $Ca_{tgt}$ values, representing different regulation for the systolic vs diastolic calcium levels. **II-A.** Calcium transient at 1000 ms pacing, illustrating how $Ca_i$ is split into systolic and diastolic values: $Ca_{i,syst}$ is defined from the beginning of the upstroke until three times the time-to-peak interval after the peak. This relative measure (instead of a single temporal threshold) is used to insure consistency at different pacing rates. $Ca_{i,dia}$ is therefore chosen to be the remainder of the CaT. The average values of $Ca_{i,syst}$ and $Ca_{i,dia}$ define the respective $Ca_{tgt}$ values, shown as dotted lines, and are 'active' (i.e., the set point of the feedback model) during the systolic or diastolic intervals, respectively. The baseline $Ca_tgt$ is shown as a solid gray line, for reference. **II-B.** The two-$Ca_{tgt}$ regulation leads to similar baseline ion channel expression. However, at 250 ms pacing, ion channel remodeling is significantly reduced. **II-C**. At 250 ms pacing, the reduced remodeling leads to a prolonged action potential and increased CaT. Note that the feedback model cannot reach either target and instead settles into an average between the two values.

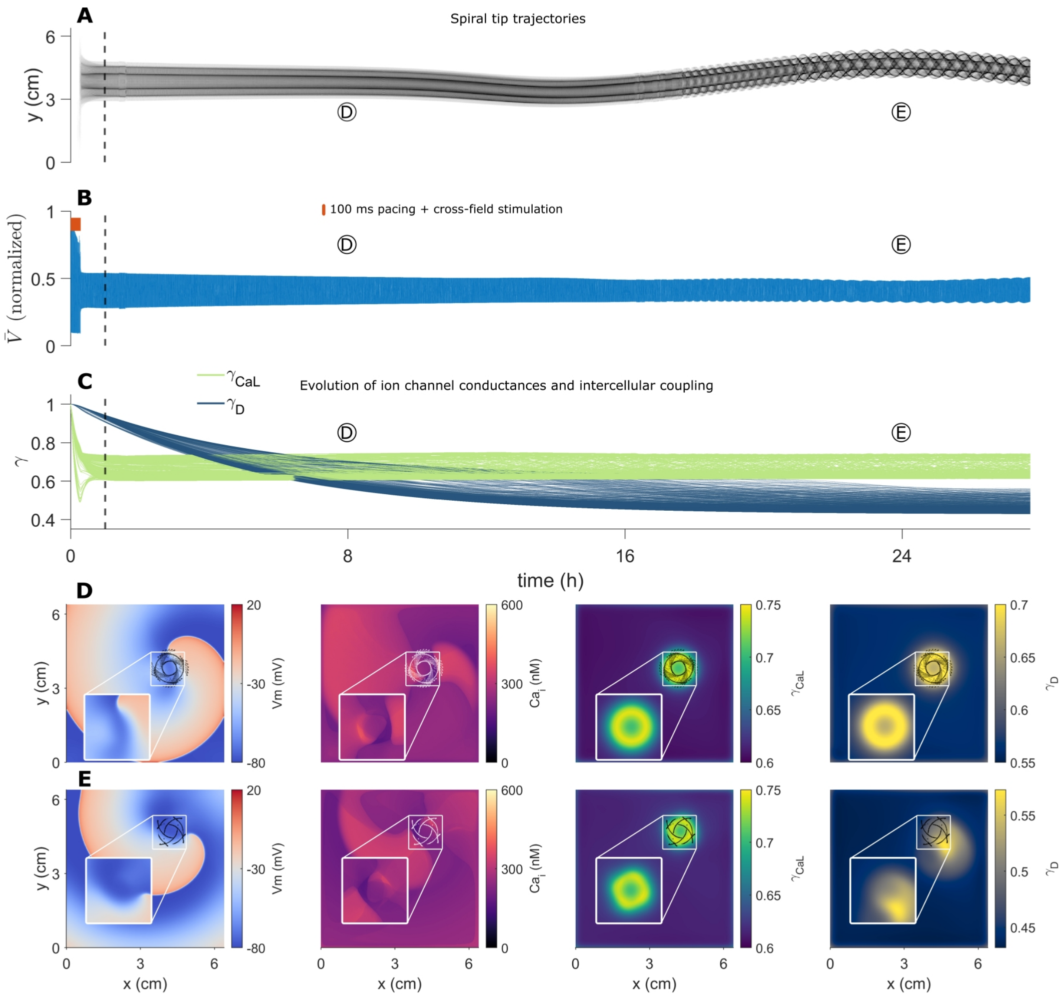

**Figure S8.** Long term spiral wave behavior in atrial tissue with homogeneous $Ca_{tgt} = 260$ nM. **A.** The position of the spiral tip trajectory (y-axis projection) and the **B.** pseudo-ECG of tissue voltage activity illustrate the initial long term stability of the spiral wave, followed by slow drift. Orange lines in (B) denote the initial rapid (100 ms) pacing followed by cross-stimulation. **C.** Long term activity illustrates the progression of the relatively faster electrical ($\gamma_{CaL}$, blue lines) and slower intercellular coupling ($\gamma_D$, green lines) remodeling. The curves denote $\gamma_{CaL}$ and $\gamma_D$ at each spatial location in the atrial tissue. Dashed vertical line in (A-C) corresponds with the time period of the simulation in Figure 2. **D, E** Snapshots showing voltage, $Ca_i$, $\gamma_{CaL}$, and $\gamma_D$ at time points corresponding to the early stable spiral (D) and the late, drifting, spiral (E). Black or white traces show spiral tip position shortly before and after each snapshot. Note the 'ring' remodeling structure apparent in both $\gamma_{CaL}$ and $\gamma_D$ (insets), corresponding to the re-entrant wave location. Simulation data is also shown in Movie S1.

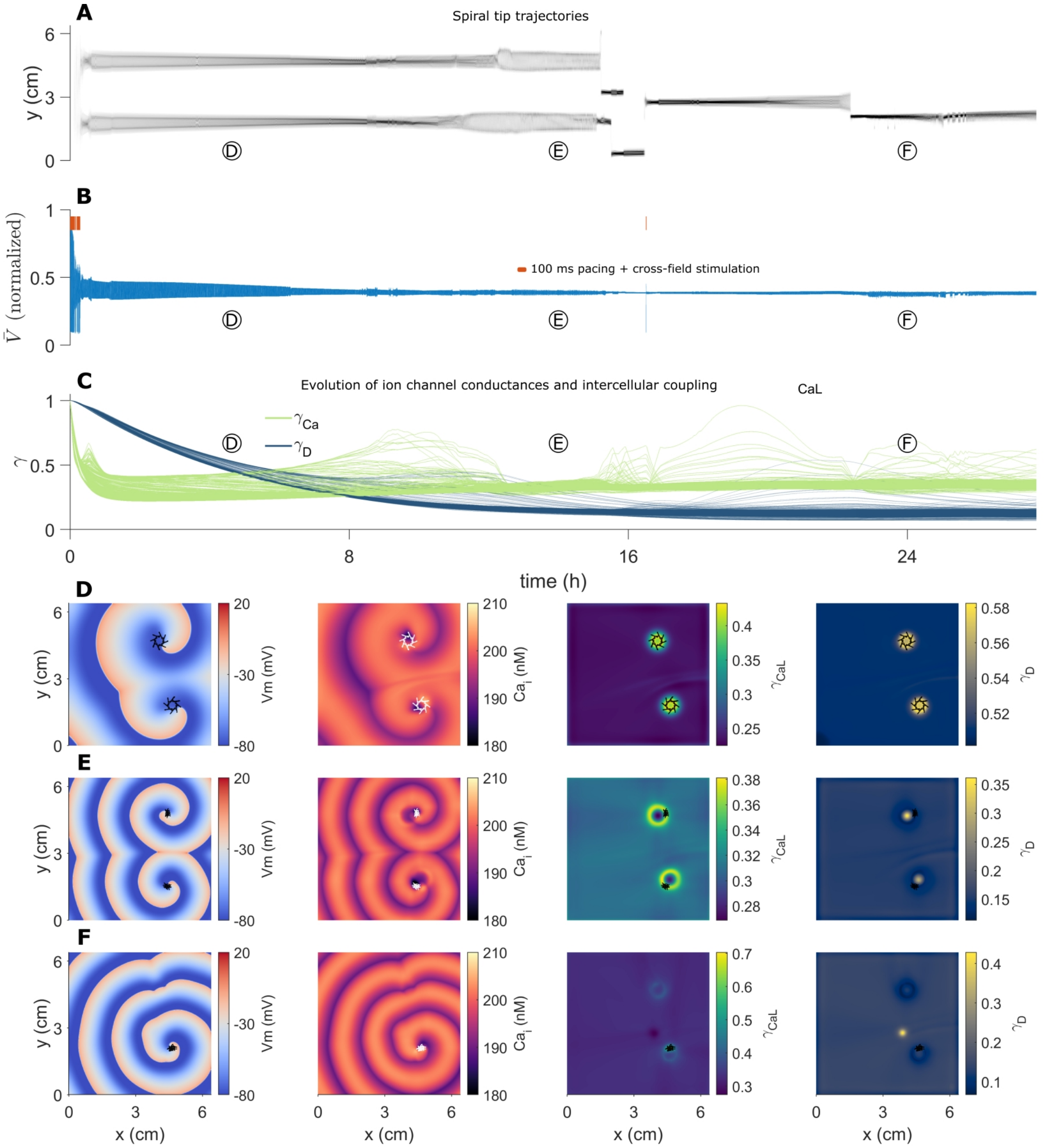

**Figure S9.** Long term spiral wave behavior in atrial tissue with homogeneous $Ca_{tgt} = 200$ nM. **A.** The position of the spiral tip trajectory (y-axis projection) and the **B.** pseudo-ECG of tissue voltage activity show significantly more dynamic activity, compared with the homogeneous tissue with $Ca_{tgt} = 258$ nM (Figure S8). Initially, two spirals are active, which then transition to a single spiral that switches locations in tissue. **C.** Progression of $\gamma_{CaL}$ and $\gamma_D$ remodeling. The overall remodeling level is more pronounced, compared with Figure S8, with a large degree of spatiotemporal variability in $\gamma_{CaL}$. **D-F.** Snapshots of transmembrane voltage, $Ca_i$, $\gamma_{CaL}$, and $\gamma_D$ at different time points. As in the single cell, the smaller $Ca_{tgt}$ leads to enhanced remodeling, and thus in atrial tissue leads to complex electrical activity. Note the complex remodeling pattern, driven by the two spiral wave spatial locations. Simulation data is also shown in Movie S2.

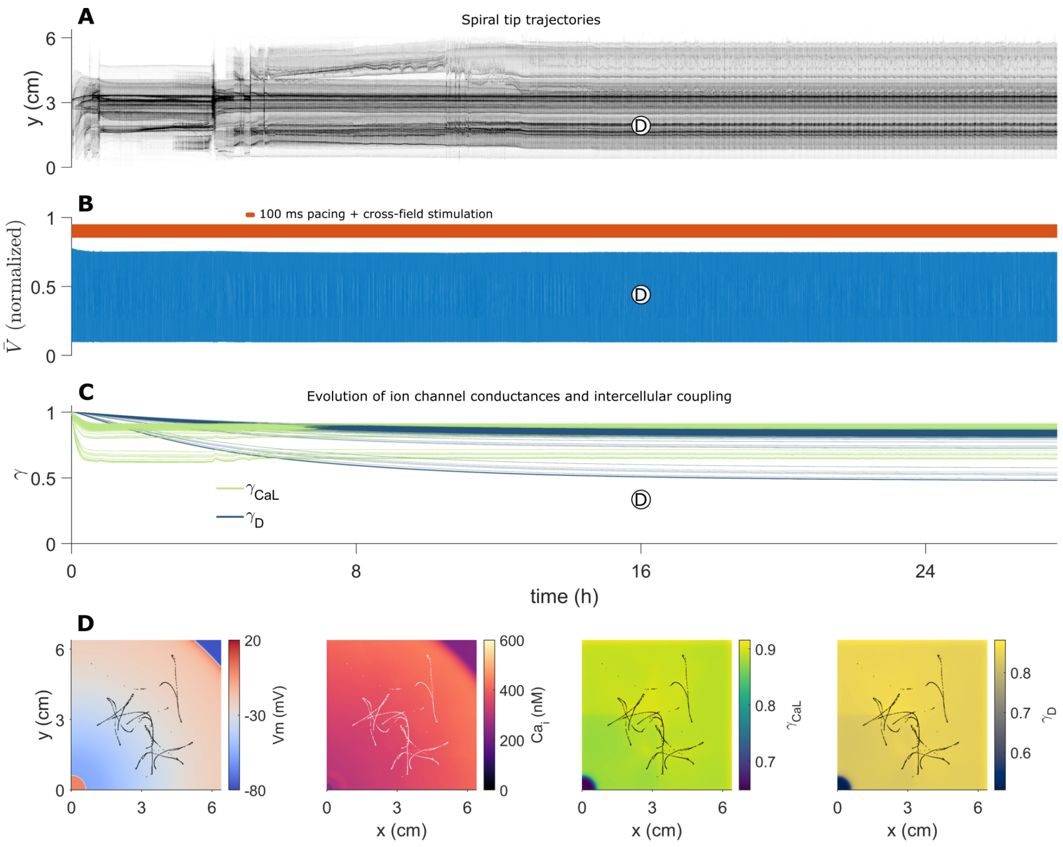

**Figure S10.** Long term re-entrant activity in atrial tissue with homogeneous $Ca_{tgt} = 320$ nM. **A.** The position of the incomplete spiral tip trajectory (y-axis projection) and the **B.** pseudo-ECG of tissue voltage. **C.** Progression of $\gamma_{CaL}$ and $\gamma_D$ remodeling. As in the single cell, the larger $Ca_{tgt}$ leads to reduced remodeling, such that in the atrial tissue, the overall degree of remodeling level is also lower, compared with than Figure S8. The lower remodeling (compared to the $Ca_{tgt} = 258$ or 200 nM cases) prevents the formation of a self-perpetuating spiral wave. **D.** Snapshots showing transmembrane voltage, $Ca_i$, $\gamma_{CaL}$, and $\gamma_D$. The waves that do form quickly extinguish. Simulation data is also shown in Movie S3.

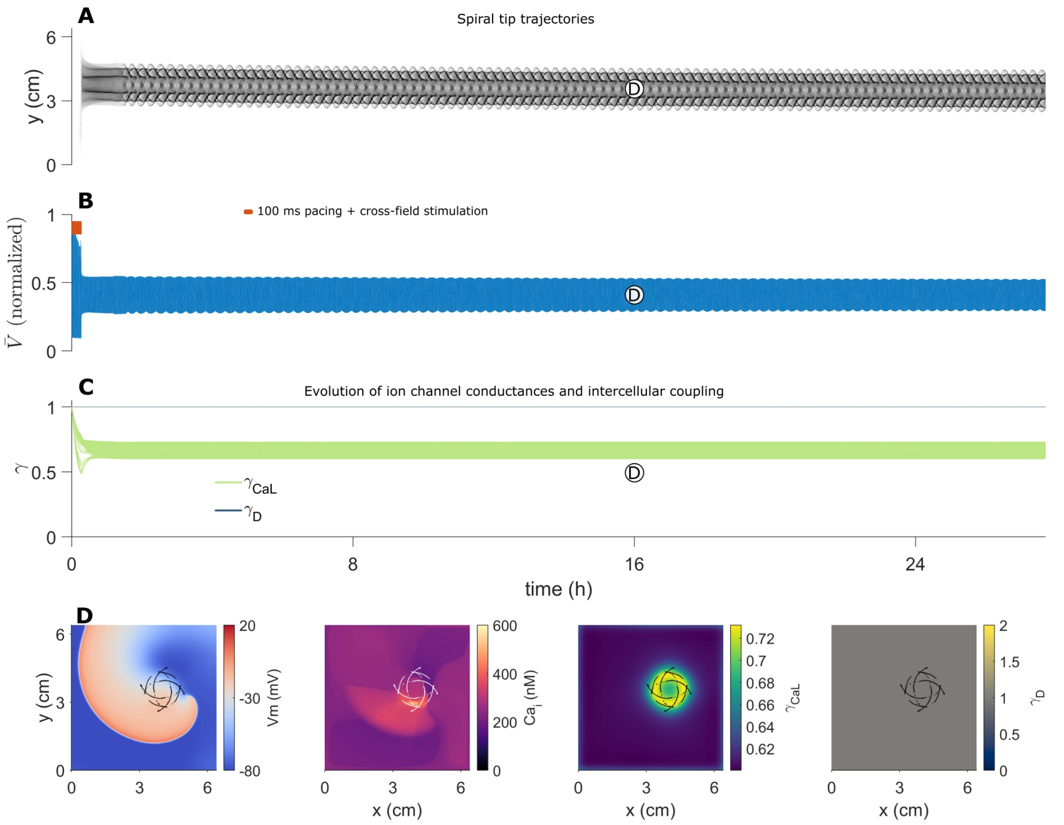

**Figure S11.** Long term re-entrant activity in atrial tissue with homogeneous $Ca_{tgt} = 258$ nM, with no intercellular coupling remodeling ($\gamma_D$ fixed at 1). **A.** The position of the spiral tip trajectory (y-axis projection) and the **B.** pseudo-ECG of tissue voltage. Note that the re-entrant wave location does not drift for the entire duration of the simulation. **C.** Progression of $\gamma_{CaL}$ remodeling. **D.** Snapshots showing transmembrane voltage, $Ca_i$, $\gamma_{CaL}$, and $\gamma_D$. Simulation data is also shown in Movie S4.

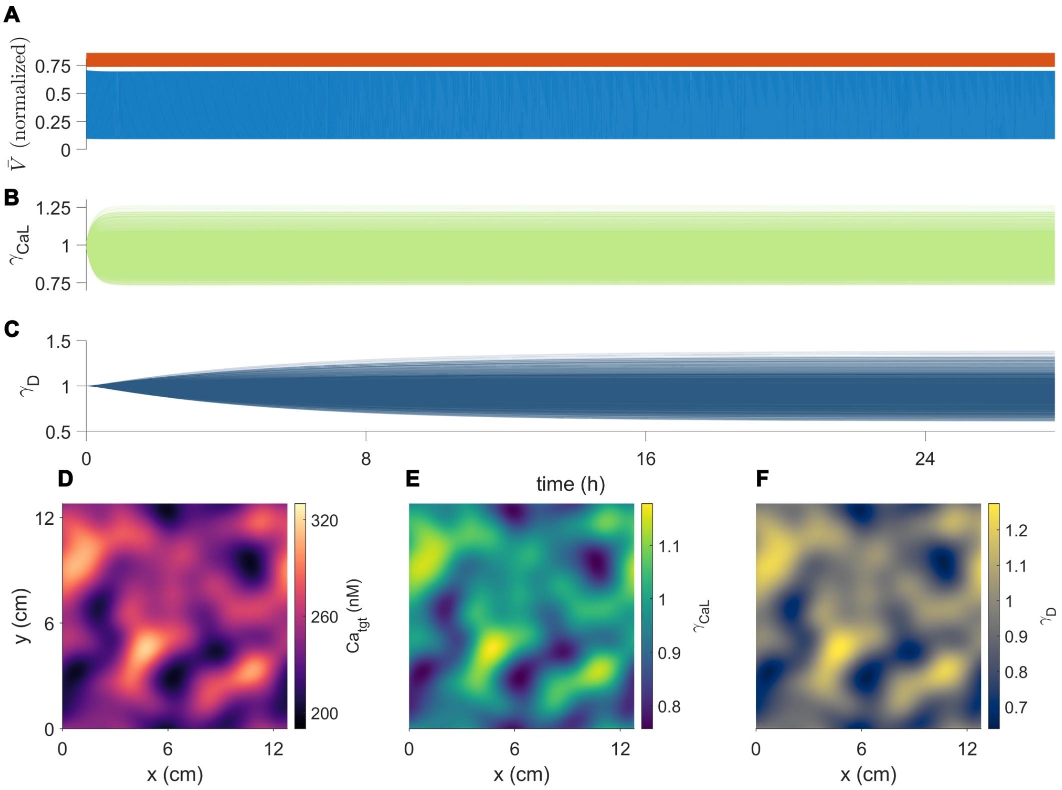

**Figure S12.** Heterogeneous 2D tissue paced at 1000 ms cycle length. **A.** Pseudo-ECG, with continuous pacing denoted by orange bar. **B.** Evolution of tissue $\gamma_{CaL}$. **C.** Evolution of $\gamma_D$. **D.** Tissue $Ca_{tgt}$ map (reproduced from Figure 3 for reference) **E.** $\gamma_{CaL}$ at steady state. **F.** $\gamma_D$ at steady state.

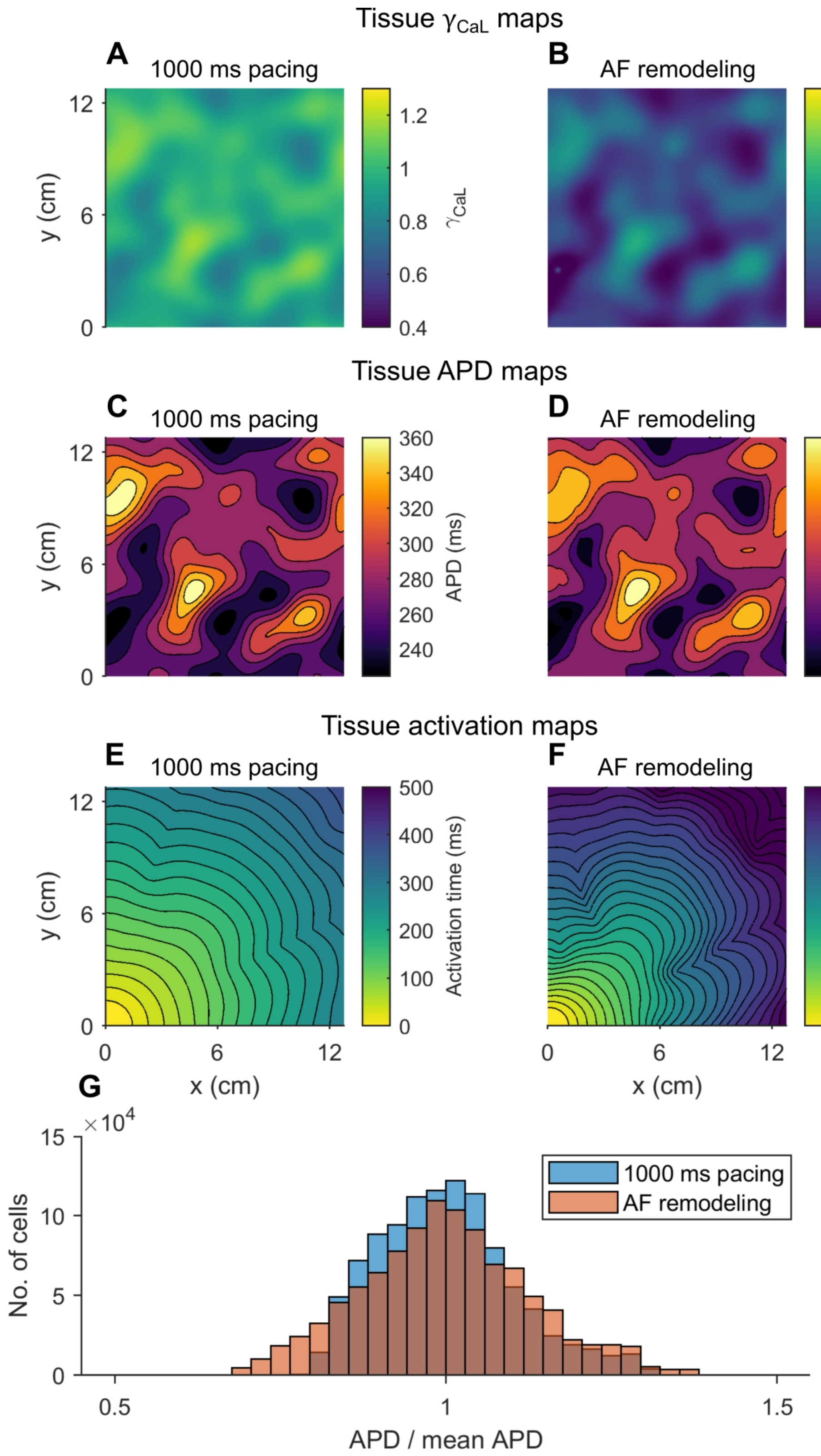

**Figure S13.** APD and conduction in healthy and AF remodeled tissue. **A, B**. Tissue $\gamma_{CaL}$ maps after long term 1000 ms pacing and atrial fibrillation (AF) remodeling **C, D**. Tissue APD map after long term 1000 ms pacing and after AF remodeling. **E, F**. Tissue activation map after long term 1000 ms pacing and after AF remodeling. Note the overall slower and irregular conduction in the AF remodeled tissue. **G.** Comparison between the APD distribution in the two cases (normalized by dividing with the mean APD value), illustrating the wider dispersion of APD values in the AF remodeled tissue.

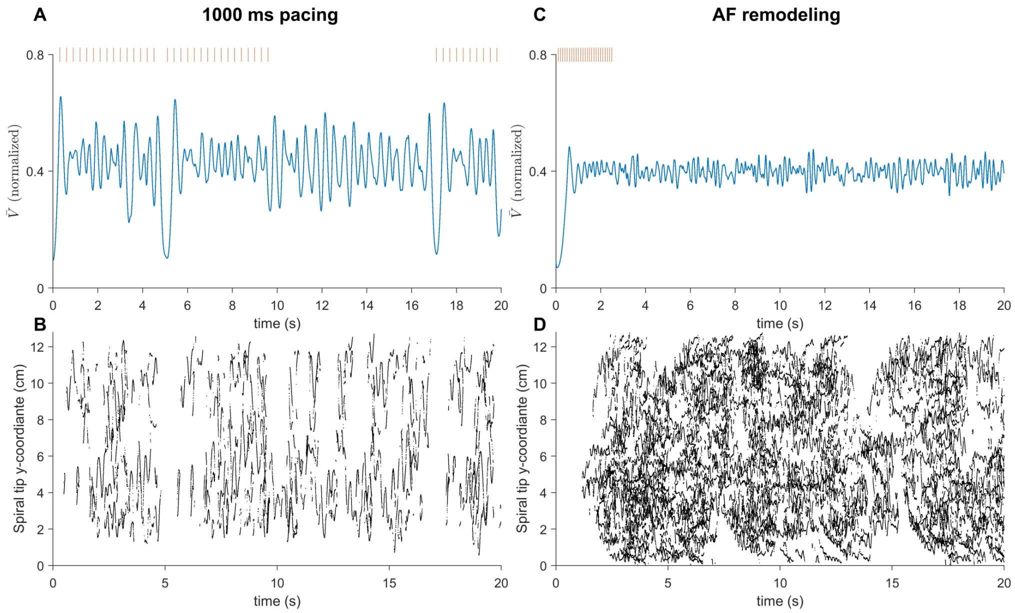

**Figure S14.** Susceptibility to re-entry in healthy and AF remodeled tissue. **A, B.** Pseudo-ECG and spiral tracks in the healthy tissue, paced at 1000 ms. Re-entry can be initiated in the healthy tissue; however episodes are brief, with few re-entrant waves. **C, D.** Pseudo-ECG and spiral tracks in the AF remodeled tissue. A single pacing episodes leads to persistent re-entry, with multiple spiral wave fibrillatory activity.

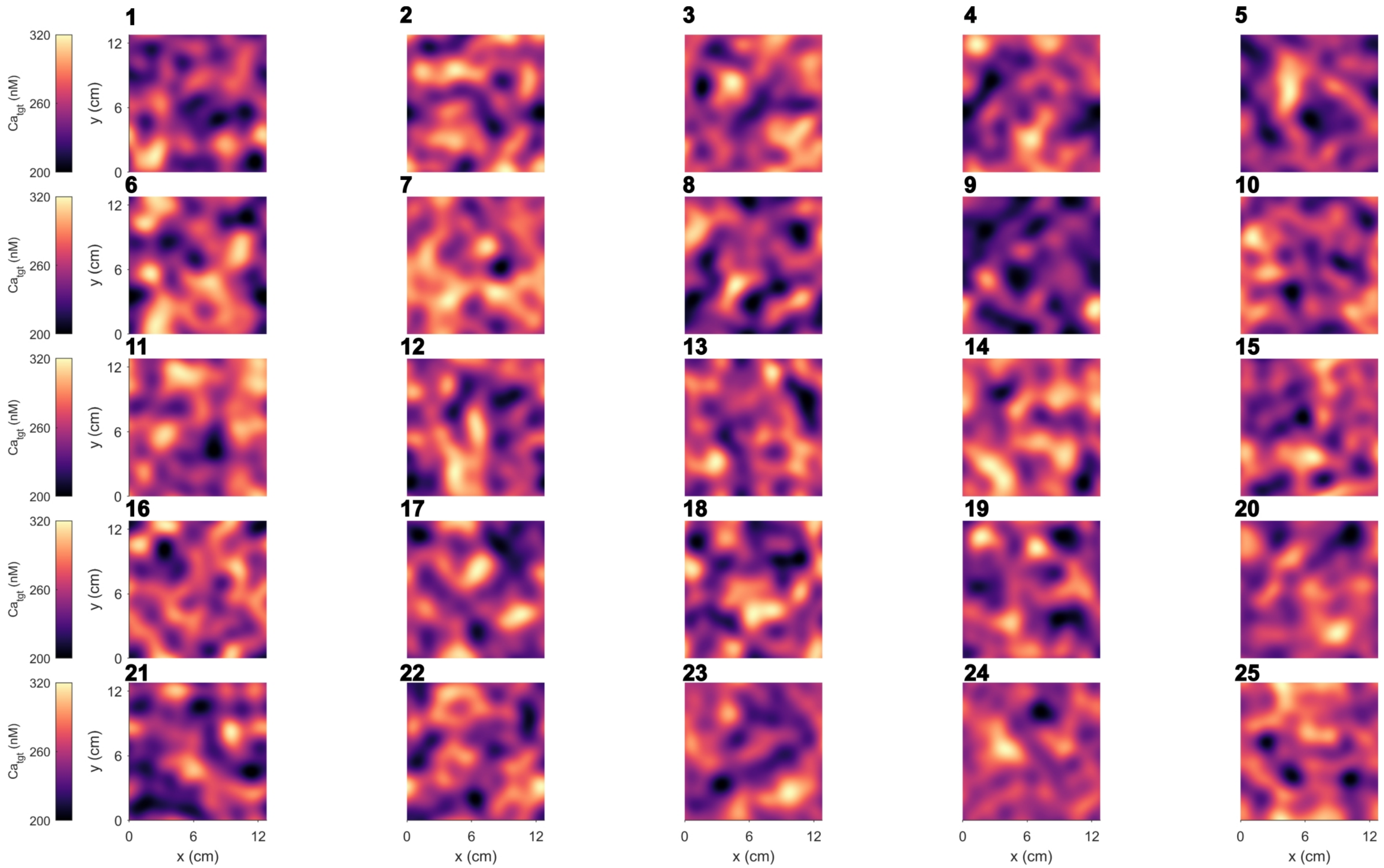

**Figure S15.** $Ca_{tgt}$ maps used for the simulations presented in Figure S16. All values are in the interval $[200 - 320]$ nM.

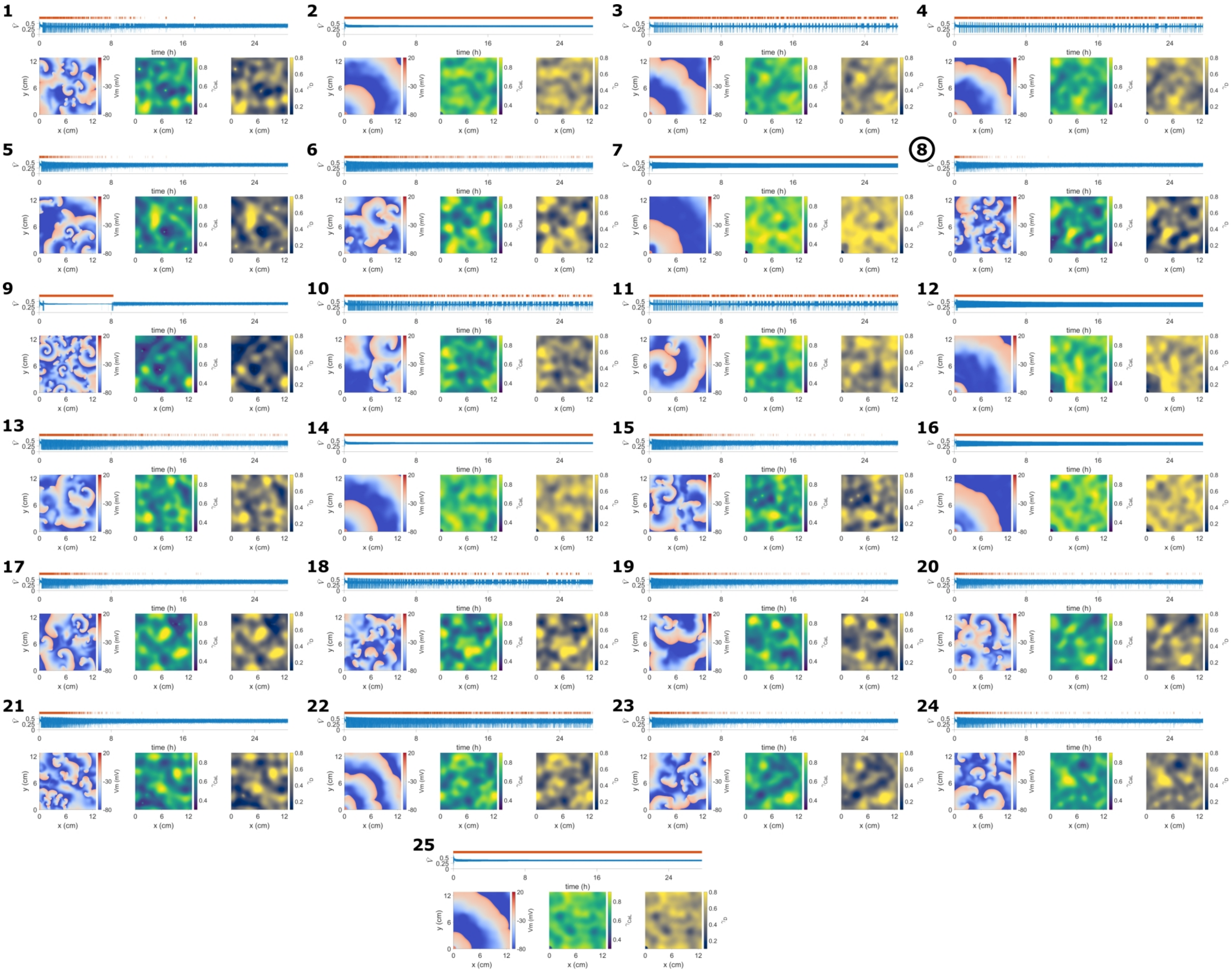
time (h)
x (cm)
y (cm)
Vm (mV)

**Figure S16.** (Previous page). Simulations on 25 different heterogeneous atrial tissues, each with different random $Ca_{tgt}$ spatial maps. Each map is a different random generation using the same map generation procedure, and each simulation follows the same protocol as in Figures 3-6. All numbered panels show (top panel) the pacing times (orange dots) and tissue pseudo-ECG, and snapshots of transmembrane voltage (left), $\gamma_{CaL}$ (middle), and $\gamma_D$ (right) at the end of simulation. The simulation highlighted in Figures 3-6 is number 8 (circled). Different $Ca_{tgt}$ spatial patterns lead to broadly 3 categories of long-term tissue dynamics: 1. no reentry is initiated, such that permanent pacing is present (2,7,12,14,16,25); 2. Intermittent re-entry and pacing (3,4,6,10,11,13,15,18,19,20,22,23,24); 3. persistent chaotic electrical activity (1,5,8,9,17,21).

### 3 Supplementary Movies

**Movie S1.** Long term spiral wave behavior in homogeneous tissue with $Ca_{tgt} = 258$ nM. The pseudo-ECG (top) spiral wave locations, in a 100 second time window for each movie frame (left), spatial maps of electrical remodeling $\gamma_{CaL}$ (middle), and intercellular coupling remodeling $\gamma_D$ (right) are shown as functions of time. *File: vid_24h_map_homog_043*

**Movie S2.** Long term spiral wave behavior in homogeneous tissue with $Ca_{tgt} = 200$ nM. The pseudo-ECG (top) spiral wave locations, in a 100 second time window for each movie frame (left), spatial maps of electrical remodeling $\gamma_{CaL}$ (middle), and intercellular coupling remodeling $\gamma_D$ (right) are shown as functions of time. *File: vid_24h_map_homog_033*

**Movie S3.** Long term spiral wave behavior in homogeneous tissue with $Ca_{tgt} = 320$ nM. The pseudo-ECG (top) spiral wave locations, in a 100 second time window for each movie frame (left), spatial maps of electrical remodeling $\gamma_{CaL}$ (middle), and intercellular coupling remodeling $\gamma_D$ (right) are shown as functions of time. *File: vid_24h_map_homog_053*

**Movie S4.** Long term spiral wave behavior in homogeneous tissue with $Ca_{tgt} = 258$ nM, with fixed $\gamma_D = 1$. The pseudo-ECG (top) spiral wave locations, in a 100 second time window for each movie frame (left), spatial maps of electrical remodeling $\gamma_{CaL}$ (middle), and intercellular coupling remodeling $\gamma_D$ (right) are shown as functions of time. *File: vid_24h_map_homog_0d*

**Movie S5.** Spiral wave initiation in heterogeneous tissue. Spatial maps of electrical remodeling $\gamma_{CaL}$ (left), transmembrane voltage $V$ (middle), and intracellular calcium $Ca_i$ (right) are shown as functions of time. This movie shows the data from Figure 4, and note that this initiation event occurs during the simulation shown in Figures 3 - 6 (map number 8, see Figures S15-S16). Recent spiral tracks are overlaid on all panels. *File: vid_init*

**Movie S6.** Spiral wave activity in remodeled heterogeneous tissue. Spatial maps of electrical remodeling $\gamma_{CaL}$ (left), transmembrane voltage $V$ (middle), and intracellular calcium $Ca_i$ (right) are shown as functions of time. This movie shows the data from Figure 6, and note that this shows activity at the end of the simulation shown in Figures 3-6(map number 8, see Figures S15-S16). Recent spiral tracks are overlaid on all panels. *File: vid_end*

**Movie S7.** Long term time-lapse spanning the entire simulation in Figures 3-6 (map 8, see Figures S15-S16). The pseudo-ECG (top) and spiral wave locations, in a 100 second time window for each movie frame (left), spatial maps of electrical remodeling $\gamma_{CaL}$ (middle), and intercellular coupling remodeling $\gamma_D$ (right) are shown as functions of time. During chaotic activity, the 'hot' spots (with high density) in the spiral location map roughly correspond to the edges of locations with increased $\gamma_{CaL}$ and $\gamma_D$. *File: vid_24h_map_8*

**Movie S8.** Long term time-lapse spanning the entire simulation in map 1, showing an example of permanent re-entry (see Figures S15-S16). The pseudo-ECG (top) and spiral wave locations, in a 100 second time window for each movie frame (left), spatial maps of electrical remodeling $\gamma_{CaL}$ (middle), and intercellular coupling remodeling $\gamma_D$ (right) are shown as functions of time. *File: vid_24h_map_1*

**Movie S9.** Long term time-lapse spanning the entire simulation in map 2, showing a tissue resistant to re-entry (see Figures S15-S16). The pseudo-ECG (top) and spiral wave locations, in a 100 second time window for each movie frame (left), spatial maps of electrical remodeling $\gamma_{CaL}$ (middle), and intercellular coupling remodeling $\gamma_D$ (right) are shown as functions of time. *File: vid_24h_map_2*

**Movie S10.** Long term time-lapse spanning the entire simulation in map 3, showing an example if intermittent re-entry and pacing (see Figures S15-S16). The pseudo-ECG (top) and spiral wave locations, in a 100 second time window for each movie frame (left), spatial maps of electrical remodeling $\gamma_{CaL}$ (middle), and intercellular coupling remodeling $\gamma_D$ (right) are shown as functions of time. *File: vid_24h_map_3*

**Movie S11.** Long term time-lapse spanning the entire simulation in map 5, showing an example of permanent re-entry (see Figures S15-S16). The pseudo-ECG (top) and spiral wave locations, in a 100 second time window for each movie frame (left), spatial maps of electrical remodeling $\gamma_{CaL}$ (middle), and intercellular coupling remodeling $\gamma_D$ (right) are shown as functions of time. *File: vid_24h_map_5*
